# Age-dependent RGS5 loss in pericytes induces cardiac dysfunction and fibrosis in the heart

**DOI:** 10.1101/2023.12.08.570774

**Authors:** Anita Tamiato, Lukas S. Tombor, Ariane Fischer, Marion Muhly-Reinholz, Leah Rebecca Vanicek, Büşra Nur Toğru, Jessica Neitz, Simone Franziska Glaser, Maximilian Merten, David Rodriguez Morales, Jeonghyeon Kwon, Stephan Klatt, Bianca Schumacher, Stefan Günther, Wesley Abplanalp, David John, Ingrid Fleming, Nina Wettschureck, Stefanie Dimmeler, Guillermo Luxán

**Author notes:** Equal contribution (Stefanie Dimmeler, Guillermo Luxán). **Corresponding authors** Guillermo Luxán Stefanie Dimmeler Goethe University Frankfurt Theodor Stern Kai 7 60590 Frankfurt; Germany.

## Abstract

**Background:** Pericytes are capillary-associated mural cells involved in the maintenance and stability of the vascular network. Although ageing is one of the main risk factors for cardiovascular disease, the consequences of ageing on cardiac pericytes are unknown.

**Methods:** In this study, we have combined single-nucleus RNA sequencing and histological analysis to determine the effects of ageing on cardiac pericytes. Furthermore, we have conducted in vivo and in vitro analysis of Regulator of G protein signalling 5 (RGS5) loss of function and finally have performed pericytes-fibroblasts co-culture studies to understand the effect of RGS5 deletion in pericytes on the neighbouring fibroblasts.

**Results:** Ageing reduced the pericyte area and capillary coverage in the murine heart. Single nucleus RNA sequencing analysis further revealed that the expression of *Rgs5* was reduced in cardiac pericytes from aged mice. In vivo and in vitro studies showed that the deletion of RGS5 impaired cardiac function, fibrosis, and induced morphological changes and a pro-fibrotic gene expression signature in pericytes characterized by the expression of different extracellular matrix components and growth factors e.g. *TGFB2* and *PDGFB.* Indeed, culturing fibroblasts with the supernatant of RGS5 deficient pericytes induced their activation as evidenced by the increased expression of α smooth muscle actin in a TGFβ2-dependent mechanism.

**Conclusions:** Our results have identified RGS5 as a crucial regulator of pericyte function during cardiac ageing. The deletion of RGS5 causes cardiac dysfunction and induces myocardial fibrosis, one of the hallmarks of cardiac ageing.

## Introduction

Alterations and functional impairment of the cardiac microvasculature are associated with heart failure and cardiac disease^1^. Ageing is a major risk factor for the development of cardiovascular disease (CVD) and critically affects both cardiac function and structure^2^. Although cardiac ageing has been well studied in cardiomyocytes, endothelial cells and fibroblasts, the molecular and cellular alterations that take place in cardiac pericytes during ageing are largely unknown.

In order to fulfil its demanding energetic tasks, the heart is a highly vascularized organ where almost every cardiomyocyte is in close contact to a capillary^3^. Pericytes are capillary-associated mesenchymal cells that ensure vascular homeostasis. They are crucial regulators of angiogenesis, vascular permeability, barrier function, and extracellular matrix formation^4^. Pericytes play a role in the mechanical support of the vasculature^5^, which is of importance in the heart, an organ constantly exposed to mechanical stress due to heartbeat^6^. Pericytes are known to have regenerative potential^7^ and there is accumulating evidence to suggest that the pericyte secretome plays an important role controlling and modifying the local environment in the heart^8^.

This study shows that ageing compromises pericyte coverage in the heart and identifies Regulator of G-protein signalling isoform 5 (RGS5) as a crucial regulator of pericyte function in the heart. RGS5 belongs to a family of proteins that act as activators of GTPases and has been described to regulate G-protein coupled receptors (GPCR) signalling, acting as an inhibitor of Gαq and Gαi^9^. In smooth muscle cells RGS5 has been shown to regulate proliferation^10^ and contraction^11^. The expression of *Rgs5* is reduced in aged cardiac pericytes and deleting RGS5 both in vivo and in vitro induces a pro-fibrotic response of these cells that, in the heart, leads to fibrosis, increased left ventricular (LV) mass, and reduced LV ejection fraction mimicking some of the features of cardiac ageing.

## Methods

### Animals

12 weeks and 18 months old male C57Bl/6J wildtype mice purchased from Janvier (Le Genest Saint-Isle, France) were used in this study.

The following transgenic mice were used: *Rgs5*^tm1a(EUCOMM)Wtsi^ (MGI ID:4432511, *Rgs5^fl/fl^*)^9^, *Pdgfrb-CreERT2*^12^. Pericyte-specific *Rgs5* knockout (*Rgs5^ΔPC^*) was generated by breeding *Rgs5^fl/fl^* mice with *Pdgfrb-CreERT2* mice. *Rgs5* deletion was induced in mice aged between 8 and 10 weeks by tamoxifen intraperitoneal injection (20 mg/ml), performed for five consecutive days (100 µl/mouse/day). Homozygous *Rgs5^fl/fl^ mice* were used for animal experiments. We compared animals carrying one copy of *Pdgfrb-CreERT2* with Cre-negative littermate controls. Experiments were performed in both females and males. The transgenic mice experiments were approved by the Regierungspraesidium Darmstadt under application number FU/2015.

### Single nucleus RNA sequencing analysis of 3-and 18 months old murine hearts

For single nucleus RNA sequencing analysis, we integrated cardiac cells from 12-week-old mice (n=3) and 18-month-old mice (n=3), as described previously^13^. We performed downstream analysis in R (4.2.1) using Seurat (4.2.1) and following the tutorials from www.satijalab.org. In brief, we used an anchor sets-based method for integration^14^ and pre-defined gene sets for cardiac cells^13,15^ to identify 10 cell types, including a cluster of pericytes (n=1792 cells) and a cluster of mixed pericyte/endothelial identity (n=113 cells). We performed differential gene expression analysis by Seurat’s *FindMarker* function using the bimodal maximum likelihood test (bimod)^16^. We considered Bonferroni adjusted p-values p < 0.05 as significant for further analysis. Gene Ontology analysis was performed using Metascape databases (https://metascape.org)^17^.

### Single nucleus RNA sequencing analysis of *Rgs5^ΔPC^* murine hearts

<u>Nuclear isolation:</u> hearts were perfused with 1X DPBS (Gibco) and directly cut into pieces. Tissue was minced in pre-filtered homogenization buffer containing 250 mM sucrose (Sigma-Aldrich), 25 mM KCl (Invitrogen), 5 mM MgCl_2_ (Invitrogen), 10 mM Tris buffer pH 8.0 (Invitrogen), 1 mM DTT (Thermo-Fisher Scientific), 1X protease inhibitor (Roche), 0.6 U/μl Ambion RNase inhibitor (Thermo-Fisher Scientific), 0.1% Triton (Sigma-Aldrich) in ultra-pure DNase/RNase-free distilled water (Thermo-Fisher Scientific). Nuclei were isolated after cell disruption with a glass dounce homogenizer (ten strokes with a loose pestle and five strokes with a tight pestle). The homogenized solution was then filtered onto a 40 μm cell strainer. The filtering was repeated onto a 20 μm cell strainer, before centrifuging the cell suspension at 500 g at 4°C for 8 min. After careful removal of the supernatant, the cell pellet was resuspended in sorting buffer containing 1% BSA (Sigma-Aldrich), 0.2 U/μl Ambion RNase inhibitor (Thermo-Fisher Scientific), 1 mM DTT (Thermo-Fisher Scientific) in DPBS (Gibco). 7AAD (Biolegend) positive nuclei were separated from cell debris using the FACSAria Fusion instrument (BD Biosciences) and sorted into collection buffer containing 1% BSA (Sigma-Aldrich), 1 U/μl Ambion RNase inhibitor (Thermo-Fisher Scientific), 1 mM DTT (Thermo-Fisher Scientific) in DPBS (Gibco). Sorted nuclei were washed twice with 0.04% BSA (Sigma-Aldrich) in DPBS (Gibco).

### Single-nucleus RNA-sequencing library preparation: sorted nuclei were subjected to single-cell

RNA sequencing using the 10X Genomics platform and the chromium Next GEM Single Cell v3.1 workflow. Briefly, 20000 nuclei were loaded on a Chip G (10X Genomics) and run in the Chromium Controller to capture and barcode poly-adenylated transcripts of single nuclei in Gel-Beads-in emulsion. Following reverse transcription, the full-length cDNA was amplified and purified with Dynabeads MyONE SILANE (10X Genomics). Single-cell gene expression libraries were obtained by enzymatic fragmentation, ligation of sequencing adapters and attachment of i5 and i7 sample indexes. A final quality control was performed on an Agilent High Sensitivity Chip using the Agilent Bioanalyzer 2000 system. Libraries were sequenced on a P2 flow cell and NovaSeq6000 instrument (Illumina) in paired-end and dual-indexing mode with an anticipated coverage of 50000 reads per nuclei.

### Single-nucleus RNA-sequencing analysis

For our initial analysis of single-nucleus RNA sequencing data, we utilized the CellRanger software suite (7.0.0) to align our data with the mouse reference genome (mm10-2020). Following the best practices for single-nucleus sequencing, we activated the *’include-introns’* option during analysis. To mitigate ambient RNA contamination, we applied the *’remove-background’* feature of CellBender (0.3.0) to the raw data counts^18^. The cleaned data was then subjected to further analysis with scanpy (1.9.6) operated within Python 3.9.18. Our downstream analysis excluded genes (expressed in less than 3 cells). We further removed cells with insufficient gene counts (fewer than 250 UMI) and with an excess of mitochondrial RNA (over 10%). We removed doublets by using *Scrublet* (0.2.3) with an estimated doublet rate of 6% and by removing the top 5% of cells with the highest number of UMI counts. After normalization of the data to total counts and a log-transformation, we identified intersections among the most variable genes using a dispersion-based approach. Dimensionality reduction and integration were achieved through principal component analysis and BBKNN batch correction approach (1.6.0.) using 3 neighboring cells within batches. Cell clustering was performed using the *Leiden* algorithm with a resolution parameter of 0.3. We visualized the data points using the UMAP technique. Clusters were annotated automatically, using *celltypist* (1.6.2., Human heart model V1.0.) and manually cross-checked with gene markers previously identified in the murine heart^13,15^.

### Echocardiography

Cardiac systolic function was assessed via echocardiography. Control and *Rgs5^ΔPC^* mice were monitored using a Vevo3100 (Fujifilm VisualSonics) before tamoxifen injection and either 4 or 8 weeks after, depending on the experimental design.

### Hypoxia assay

Hypoxic areas in cardiac tissue were assessed in Control and *Rgs5^ΔPC^* mice using Hypoxiprobe Plus kit-FITC (HP2-100, Hypoxiprobe). Animals were injected intraperitoneally with 60 mg/ml of pimonidazole 20 minutes before sacrifize. Paraffin sections were stained for Primary FITC-Mab1 (1:50, HP2-100, Hypoxiprobe) and anti-FITC HRP linked antibody (1:100, HP2-100, Hypoxiprobe) following manufacturer’s instructions. Signal was amplified using 3,3′-Diaminobenzidine (DAB, 1856090, Thermo Fisher). Nuclei were identified using Eosin staining. A positive hypoxic femur section was included as a positive experimental control.

### Genomic DNA extraction and PCR

To check whether the tamoxifen-induced deletion was successful, total genomic DNA was extracted from the apex of controls and *Rgs5^ΔPC^* mice using the DNeasy Blood and Tissue Kit (69504, Qiagen) and PCR was performed using KOD Xtreme™ HotStart DNA Polymerase Kit (71975-3, Novagen) following manufacturer’s instructions. We used primers flanking exon 3 (forward: CTGCCAGCCTGCTTTAATTGG, reverse: TGAAGCTGGCAAATCCATAGC).

### Immunohistochemistry of murine hearts

Murine hearts were perfused and fixed with 4% PFA (28908, Thermo Fisher) for 3 hours and subsequently washed with PBS. The hearts were then dehydrated in a sucrose (S0389, Sigma-Aldrich) gradient (5% 2 hours, 10% 2 hours and 20% overnight) at 4°C and embedded in a 15% sucrose, 8% gelatine (G1890, Sigma-Aldrich) and 1% polyvinylpyrrolidone (PVP, P5288, Sigma-Aldrich) solution. The hearts were sectioned 50 μm thick using a Cryostat (CM3050, Leica) and stored at −20°C before immunofluorescence processing. After re-hydration, sections were permeabilised using 0.3% Triton X-100 (3051.2, Roth) in PBS 3 times, 10 minutes each and blocked in 3% bovine serum albumin (BSA, A7030, Sigma-Aldrich), 0.1% Triton X-100, 20 mM MgCl_2_ (2189.2, Roth) and 5% donkey serum (ab7475, Abcam) in PBS for 1 hour at room temperature. Primary antibodies were incubated overnight at 4°C in a humidity box. Primary antibodies: rat anti-PDGFRβ (1:50, 14-1402-82, Invitrogen), rabbit anti-NG2 (1:100, AB5320, Millipore), rat anti-CD45 (1:50, 103102, Biolegend), rabbit anti-CD68 (1:100, 97778S, Cell Signalling), rabbit anti-Albumin (1:100, ab19196, Abcam), Isolectin B4 biotinylated (1:25, B-1205, Vector), goat anti-PDGFRα (1:100, AF1062, R&D), rabbit anti-RGS5 (1:100, PA5-75560, Invitrogen). After 3 PBS washes of 10 minutes each, secondary antibodies and DAPI (D9542, Sigma-Aldrich) were incubated for 1 hour at room temperature in PBS 5% BSA. Secondary antibodies: donkey anti-rat Alexa Fluor™ 594 (1:200, A21209, Invitrogen), donkey anti-rabbit Alexa Fluor™ 647 (1:200, A31573, Invitrogen), Streptavidin Alexa Fluor™ 555 conjugate (1:200, S32355, Invitrogen), donkey anti-goat Alexa Fluor® 488 (1:200, A11055, Invitrogen), wheat germ agglutinin (WGA) 647 conjugate (1:300, W32466, Life Technologies). Slides were mounted with Fluoromount-G™ Mounting Medium (4958-02, Thermo-Fisher) and immunofluorescence images were acquired using a Leica Stellaris confocal microscope.

### Quantitative analysis of pericytes

All pericyte quantifications were done on high-resolution confocal images of thick z-sections of the heart using Volocity (x64) software (version 6.5.1, Quorum Technologies, Canada). Pictures acquisition was performed using a 63X immersion objective. The pericyte coverage was determined by defining the intersection of pericyte volume (PDGFRβ^+^ NG2^+^) and endothelial cells (isolectinB4^+^) volume. We considered pericyte-cell bodies all nuclei surrounded by pericyte markers (PDGFRβ^+^ NG2^+^ DAPI^+^). We defined pericyte branches as all the PDGFRβ^+^ NG2^+^ protrusions departing from pericyte cell bodies. All values were quantified from a minimum of six left ventricle areas per sample and were finally normalised to the total isolectinB4 volume.

### Collagen deposition analysis

To analyse collagen deposition in the heart, we performed Sirius Red staining on paraffin sections. Hearts fixed as described above were embedded in paraffin. After deparaffinization and rehydration, the slides were stained with 0.1% Picro Sirius Red solution prepared using Sirius Red F3BA (1A280, Waldeck GmbH) in an aqueous solution of Picric Acid (6744, Sigma Aldrich). Slides were mounted with Pertex mounting medium (00801-EX, HistoLab) and images were acquired using a Nikon Eclipse Ci microscope with a 40X objective. Positive area was measured using ImageJ (NIH, version 1.53a) and normalised to the total tissue area. Perivascular fibrosis was normalized to the calibre of the vessel.

### β-Galactosidase staining

Senescence staining was performed on paraffin sections using using the β-galactosidase staining kit (9860S, Cell Signaling) following the manufacturer’s instructions. Images were acquired using a Nikon Eclipse Ci microscope with a 40X objective.

### Cell culture

Human Placenta Pericytes (hPC-PL, C-12980, PromoCell) were seeded at 6000 cells/cm^2^ density and culture in the respective Pericyte Growth Medium 2 (C-28041, PromoCell). Upon reaching 90% confluency, cells were washed with medium and detached by enzymatic digestion using Accutase (A6964, Sigma-Aldrich) (37°C/5% CO_2_; 3 minutes). We used hPC-PL from different donors (464Z005, 463Z019.1, 422Z033.3), both female and male. All experiments were performed between passage 2 and 6.

Human Brain Vascular Pericytes (HBVP, 1200, 27194, ScienCell) were cultured on 1% Gelatin (G1393, Merck) in DMEM Glutamax (31966-021, Gibco) supplemented with 10% Fetal bovine serum (10270106, Gibco) and 1% Pen/Strep (11074440001, Roche).

Human Umbilical Vein Endothelial Cells (HUVEC, CC-2519, Lonza) were cultured in Endothelial Basal Medium (EBM, CC-3121, Lonza) supplemented with 10% FBS (10270106, Gibco), Amphotericin-B, ascorbic acid, bovine brain extract, endothelial growth factor gentamycin sulphate, and hydrocortisone (EGM SingleQuots Kit, CC-4133, Lonza). HUVECs were used up to passage 4.

Human cardiac fibroblasts (HCF, C-12375, Promocell, 424Z011.9 and 437Z012.4) were seeded at 4000 cells/cm^2^ density and cultured in the respective Fibroblast Growth Medium (C-23025, PromoCell).

### Gene silencing and cell treatments

*RGS5* silencing was performed using siRNA. *RGS5* siRNA (ID: SASI_Hs01_00094781, Sigma-Aldrich) and control siRNA (sense: CGUACGCGGAAUACUUCGA; antisense: UCGAAGUAUUCCGCGUACG, Sigma-Aldrich)^19^ were used at a final concentration of 50nM. In hBVPs, *RGS5* silencing was performed using MISSION® esiRNA (EHU042281, Sigma-Aldrich) at 67nM final concentration. hPC-PL were transfected using Lipofectamine™ RNAiMAX Transfection Reagent (13778150, Thermo-Fischer Scientific) for 4 hours in Optimem (51985034, Gibco). RNA or protein isolation was performed 48 hours after transfection. hPC-PL were treated with the Gαq agonist U-46619 (10 μM, 1h, 37°C, 16450, Cayman). U-46619 is a thromboxane A_2_ receptor agonist, that signals via the Gαq-mediated cascade. hPC-PL supernatant from *siControl* and *siRGS5* samples was applied on HCF for 48 hours and TGFβ signal neutralized using anti TGFβ 1,2,3 monoclonal antibody (1mg/ml, Thermo-Fisher Scientific) and control IgG antibody (1mg/ml, Thermo-Fisher Scientific).

### IP1 production assay

Inositol 1 phosphate (IP1) accumulation in hPC-PL was measured using the HTRF IP-One Gαq Detection Kit (Cisbio). This assay enables the direct pharmacological activity of Gαq - coupled receptors. Upon *RGS5* knockdown, hPC-PL were detached with accutase and IP production was measured following manufacturer’s instructions.

### Immunocytochemistry of human cardiac fibroblasts

HCF were cultured in μ-Slide 4 Well (80426, Ibidi) and treated with hPC-PL supernatant from *siControl* and *siRGS5* samples for 48 hours. After washing with PBS, cells where fixed in 4% PFA for 10 min and subsequently washed with PBS (3x, 5 minutes). Cells were then permeabilised using 0.1% Triton X-100 in PBS for 10 minutes and blocked in 5% donkey serum in PBS for 1 hour at room temperature. Primary antibodies were incubated overnight at 4°C in a humidity box. Primary antibodies: anti-Actin α Smooth Muscle - Cy3™ (1:300, C6198, Sigma-Aldrich), Oregon green™ 488 phalloidin (07466, Invitrogen). After 3 PBS washes of 5 minutes each, DAPI was incubated for 15 minutes at room temperature in PBS. Cells were mounted with Fluoromount-G™ Mounting Medium (4958-02, Thermo-Fisher) and immunofluorescence images were acquired using a Leica Stellaris confocal microscope.

### Proliferation assay

Cell cycle phases were analysed using the BrdU Flow kit (51-2354AK, BD Biosciences). hPC-PL were treated with 10 µM BrdU in culture medium for 4 hours at 37°C. Cell pellets were then processed according to the manufacturer’s instructions. In brief, after permeabilization, cells were incubated with 300 µg/ml DNase I for 1 hour at 37°C, washed with Perm/Wash Buffer and stained with V450 anti-BrdU antibody (10 µg/mL, 560810, BD Biosciences) for 20 minutes at room temperature. Finally, 7-AAD (1 µg/mL, 555816, BD Biosciences) was added for 10 minutes at room temperature in dark and cells were analysed on a LSRFortessa X-20 flow cytometer (BD Biosciences) and FlowJo Software 10.8.1.

### Migration assay

To assess the migratory capacity of hPC-PL upon *RGS5* knockdown, transfected cells were cultured in 2-well inserts (80209, Ibidi) at 25000 cells/well density for 24 hours. After removal of the insert, cells were washed, and images of the cell-free gap were taken using an Eclipse Ti2 inverted microscope (Nikon) at time 0 and again after 10 hours. The cell-free area was measured using the software ImageJ (version 1.52p) and migration capacity was calculated as % of closed area from time 0.

### Pericyte-endothelial cell adhesion assay

The interaction between pericytes and endothelial cells was assessed using a 3D Matrigel Matrix (#354230, Corning). To allow imaging, the experiment was performed using viral transfected GFP-expressing HBVPs^20,21^, which were subsequently transfected with siRNA as described above, while HUVECs were seeded in medium supplemented with Red-Dil-Ac-LDL (10 µg/mL, L3484, Invitrogen) and incubated overnight at 37°C. In brief, HUVECs were seeded on top of the gel and incubated for 3 hours at 37°C. GFP-expressing transfected HBVPs were seeded on top and incubated for further 3 hours at 37°C. Afterwards, the medium was removed and another layer of Matrigel was added and incubated for 30 minutes at 37°C. A customized 1:1 ratio medium made of HBVPs medium and HUVEC medium in equal parts was added to the gel and cells were incubated overnight. After fixation in ROTI® Histofix 4% (P087.4, Carl Roth) at 4°C in the Matrigel for 30 minutes, images were acquired 72 hours after transfection with a Leica SP8 confocal microscope. Analysis was performed by using Volocity (x64) software (version 6.5.1, Quorum Technologies, Canada). The total number of HBVPs per image was counted and subsequently, the number of HBVPs with compromised morphology. The number of HBVPs that were completely spherical was not included in the quantification. The number of morphologically compromised HBVPs was normalized to the total area of Dil[1]Ac-LDL stained HUVECs.

### RNA isolation, cDNA synthesis and quantitative RT-qPCR

Total RNA was extracted from cells using the RNeasy Plus Mini Kit (Qiagen) according to the manufacturer’s instructions. The synthesis of complementary DNA (cDNA) was performed by the reverse transcription (RT) of mRNA using M-MLV reverse transcriptase (Thermo-Fisher Scientific). Quantitative PCR reactions were performed using the StepOnePlus real-time PCR cycler (Thermo-Fisher Scientific) and relative gene expression was calculated by normalisation to *ACTB* gene expression (2^-ΔCt^). Human primers used: *RGS5* (f: AGAAACCAGCCAAGACCCA, r: GGAGTTTGTCCAGGGAATCAC), *ACTB* (f: CATGTACGTTGCTATCCAGGC, r: CTCCTTAATGTCACGCACGAT), *ACTA2* (f: CATCATGCGTCTGGATCTGG, r: GACAATCTCACGCTCAGCAG), *TGFB2* (f: AAGAAGCGTGCTTTGGATGCGG, r: ATGCTCCAGCACAGAAGTTGGC).

### Bulk RNA sequencing

Total RNA was used as input for whole RNA-Seq library preparation, for which integrity was verified with LabChip Gx Touch 24 (CLS138162, Perkin Elmer). 1 µg of total RNA was used as input for Truseq Stranded total RNA library preparation following the low sample protocol (Illumina). Sequencing was performed on the NextSeq2000 instrument (Illumina) using P3 flowcell with 1x72bp single end setup. The resulting raw reads were assessed for quality, adapter content and duplication rates with FastQC (http://www.bioinformatics.babraham.ac.uk/projects/fastqc). Trimmomatic version 0.39 was employed to trim reads after a quality drop below a mean of Q20 in a window of 5 nucleotides^22^. Only reads between 30 and 150 nucleotides were cleared for further analyses. Trimmed and filtered reads were aligned versus the Ensembl human genome version hg38 (Ensembl release 101) using STAR 2.7.9a with the parameter “-- outFilterMismatchNoverLmax 0.1” to increase the maximum ratio of mismatches to mapped length to 10%^23^. The number of reads aligning to genes was counted with featureCounts 2.0.2. tool from the Subread package^24^. Only reads mapping at least partially inside exons were admitted and aggregated per gene. Reads overlapping multiple genes or aligning to multiple regions were excluded. A combined raw count matrix was calculated, and batch corrected per data set using CountClust^25^. The batch-corrected matrix was used for differential expression analysis using DESeq2 version 1.30.0^26^. Only genes with a minimum fold change of ±2 (log2 = ±1), a maximum Benjamini-Hochberg corrected p value of 0.05, and a minimum combined mean of 5 reads were deemed to be significantly differentially expressed. The Ensemble annotation was enriched with UniProt data (release 17.12.2018) based on Ensembl gene identifiers (Activities at the Universal Protein Resource (UniProt)). Gene Ontology analysis was performed using the Enrichr, BioPlanet 2019 (https://maayanlab.cloud/Enrichr/)^27–29^.

### Cell lysis, protein isolation and western blot

Apex from Control and *Rgs5^ΔPC^* hearts were homogenized and lysed in RIPA buffer (R0278, Sigma-Aldrich) supplemented with protease (1:50, P8340, Sigma-Aldrich) and phosphatase inhibitors (1:100, P5726, Sigma-Aldrich). Protein concentration was measured using the DC Protein Assay Kit II (5000112, Bio-Rad) following manufacturer’s instructions. After denaturation at 95°C, proteins were loaded on pre-casted gradient gels (4561094/5678094, Bio-Rad) and separated by SDS-PAGE in 1X Tris/Glycine/SDS Buffer (1610732, Bio-Rad). Transfer onto nitrocellulose membrane (10600006, Cytiva) was performed in 1X Tris/Glycine Buffer (1610734, Bio-Rad) using the Trans-Blot® Semi-Dry system (1703940, Bio-Rad). Membranes were blocked in 5% milk (sc-2324, Santa Cruz) and probed with primary antibodies overnight at 4°C followed by incubation with ECL-HRP-linked secondary antibody (1:1000, NA934V, Amersham). Primary antibodies used: rabbit anti-GAPDH (1:1000, 2118S, Cell Signalling), rabbit anti-RGS5 (1:500, ab-196799, Abcam). Proteins were detected by chemiluminescence using HRP Substrate (WBKLS0500, Millipore) in a ChemiDoc Touch imaging system (Bio-Rad). Bands were quantified by densitometry with ImageJ (version 1.52p).

### Metabolomics

#### Pericyte Extraction

Trifluoroethanol:H_2_O (1:1, 100 µl) was added to samples before vortexing and incubation on ice for 10 minutes. Next, 200 µl of methanol:ethanol (1:1) was added. Samples were again vortexed and left on ice for 10 minutes. Next, 200 µl of ice-cold H_2_O containing the standards;^13^C-fumarate-C_2_, ^13^C-citrate-C_2_ and ^13^C-succinate-C_1,4_ at a concentration of 0.02 mM was added. After 10 min, samples were sonicated on ice for 30 sec, vortexed, and centrifuged (14 000 rpm, 10 minutes, 4 °C). The supernatant was transferred to two tubes and the solvent fraction was evaporated under a stream of N_2_-gas. The remaining aqueous fraction was frozen at − 80 °C and freeze-dried using the Alpha 3-4 LSCbasic system (Martin Christ, Osterode am Harz, Germany). Dried samples were reconstituted in H_2_O:AcN (1:3; tube 1 for analysis of redox metabolites & nucleotides/nucleosides), or in 100 % H_2_O containing 0.2 % formic acid (tube 2 for analysis of TCA cycle metabolites). Samples were centrifuged (14 000 rpm, 10 minutes, 4 °C) and the supernatant was transferred to MS vials and subjected to mass spectrometric analysis.

#### Quantification of metabolites via liquid-chromatography mass spectrometry (LC-MS)

*Redox metabolites, nucleotides & nucleosides*: Samples (4 µL) were injected via an Infinity II Bio liquid chromatography system into a 6495C triple quadrupole mass spectrometer (both Agilent Technologies, Waldbronn/Germany). Metabolites were separated on an Acquity BEH Amide HILIC column (1.7 µm, 2.1 x 100 mm, Waters) by using the following mobile phase binary solvent system and gradient at a flow-rate of 0.4 mL/min: Mobile A consisted of 100 % water with 10 mM ammonium acetate and 5 µM medronic acid at a pH of 9.3. Mobile phase B consisted of 100 % acetonitrile with 10 mM ammonium acetate. The following 23 min gradient program was used: 0 min 85 % B, 0-1 min 85 % B, 1-8 min 75 % B, 8-12 min 60 % B, 12-15 min 10 % B, 15-18 min 10 % B, 18-19 min 85 % B, 19-23 min 85 % B. The column compartment was set to 20 °C. Metabolites were detected with authentic standards and/or via their accurate mass, fragmentation pattern and retention time in polarity switching ionization dynamic MRM AJS-ESI mode, and quantified (where appropriate) via a calibration curve. The gas temperature of the mass spectrometer was set to 200 °C and the gas flow to 14 L/min. The nebulizer was set to 40 psi. The sheath gas flow was set to 12 L/min, with a temperature of 375 °C. The capillary voltages were set at 4000/2500 V with a nozzle voltage of 500/0 V. The voltages of the High-Pressure RF and Low-Pressure RF were set to 150/90 and 60/60 V, respectively. Metabolite peaks were annotated with Skyline-daily (version 22.2.1.278).

#### TCA cycle metabolites

Samples (8 µL) were injected as outlined above. Metabolites were separated on an Acquity HSS T3 C18 column (1.8 µm, 2.1 x 150 mm, Waters) by using the following mobile phase binary solvent system and gradient at a flow-rate of 0.35 mL/min: Mobile A consisted of 100 % water with 0.2 % formic acid. Mobile phase B consisted of 100 % acetonitrile with 0.2 % formic acid. The following 11 min gradient program was used: 0 min 1 % B, 0-6 min 1 % B, 6-7 min 80 % B, 7-8 min 80 % B, 8-11 min 1 % B. The column compartment was set to 30 °C. Metabolites were detected as mentioned above. The gas temperature of the mass spectrometer was set to 240 °C and the gas flow to 19 L/min. The nebulizer was set to 50 psi. The sheath gas flow was set to 11 L/min, with a temperature of 400 °C. The capillary voltages were set at 1000/1000 V with a nozzle voltage of 500/500 V. The voltages of the High-Pressure RF and Low-Pressure RF were set to 100/100 and 70/70 V, respectively. Metabolite peaks were annotated with Skyline-daily (version 22.2.1.278).

#### Statistics

Statistical analyses were performed using Prism 9.2.0 (GraphPad Software Inc.). Shapiro-Wilk test was used to assess the normal distribution of values. For normally distributed data, unpaired, two-tailed Student’s t-test was used. Non-parametric data were analysed with Mann-Whitney test. For multiple comparisons, ordinary one-way ANOVA test followed by Tukey’s multiple comparisons test was used. Data are represented as mean ± SEM and a P value < 0.05 was considered significant.

## Results

### *Rgs5* expression is reduced in the old heart

In order to study the effect of ageing on the heart, we performed a sequential histological analysis of the heart at different stages of ageing. We observed a continous increase in cardiac fibrosis, that was significant from 16 months of age onward (**Supplementary** Figure 1A**, B**). This increase in fibrosis is followed by hypertrophy. Cardiomyocyte relative area is significantly increased from 18 months of age (**Supplementary** Figure 1C**, D**). Ageing is characterized by a reduction of microvascular density in the myocardium^30^. Histological analysis of cardiac pericytes revealed that the total pericyte area was reduced with ageing and becomes significant in 22 month-old mice (**Supplementary** Figure 1E**, F**). Interestingly, pericyte coverage, which is defined as the pericyte area located in contact with capillaries, is reduced from 18 months of age (**Figure 1A-D**) accompanied by a clear change in pericyte morphology. Pericytes of the aged heart presented a disorganized phenotype with wider cellular extensions that did not aligned with the capillaries (**Supplementary** Figure 1G, arrowheads). Furthermore, despite the reduction of pericyte area, we observed increased non-pericyte PDGFRβ area in the 18-month-old mice hearts (**Supplementary** Figure 1G**, H**). These data suggest that a decreased pericyte coverage of cardiac capillaries coincides with other hallmarks of ageing.

**Figure 1.**
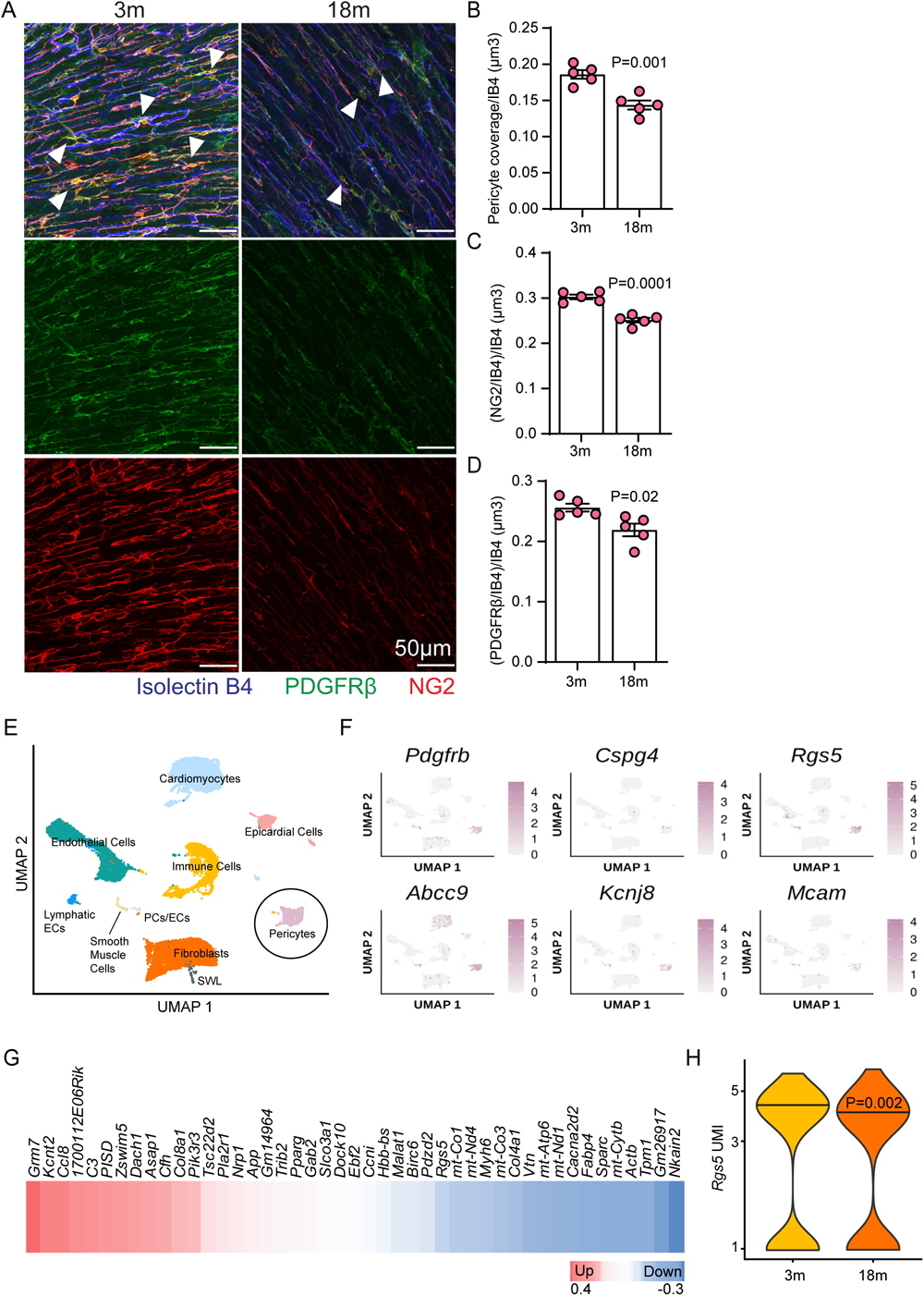
Pericytes are reduced in the old heart and *Rgs5* expression is downregulated in the aged heart. **A.** Immunofluorescence staining of left ventricular cross sections of 3 and 18 months old C57BL/6 murine hearts. Pericytes are identified as double PDGFRβ and NG2 positive cells (arrowheads). **B.** Quantification of pericyte coverage normalized to the vasculature area. **C.** Quantification of NG2 coverage normalized to the vasculature area. **D.** Quantification of PDGFRβ coverage normalized to the vasculature area. **B-D.** n=5 per condition. Data are shown as mean ± SEM. P value was calculated using two tailed unpaired *t*-test. **E.** Uniform Manifold Approximation and Projection (UMAP) plot showing cell-type specific clustering of all data points from cardiac single-nuclei sequencing. Across both 3 months old (n=3; 14247 cells) and 18 months old (n=3; 12402 cells) samples, we identified ten individual cell types: Cardiomyocytes, Endothelial Cells, Epicardial Cells, Fibroblasts, Immune Cells, Lymphatic Endothelial Cells (ECs), Pericytes, Pericyte/Endothelial mixed cluster (PCs/ECs), Schwann-like Cells (SWL), Smooth Muscle Cells. **F.** Feature plot showing gene expression of pericyte marker genes (*Pdgfrb, Cspg4, Rgs5, Abcc9, Kcnj8, Mcam*). The colored scale bar indicates the log-normalized gene expression level. **G.** Heatmap showing differentially expressed genes (DEG) between 3 and 18 months old pericytes. Upregulated genes are represented in red and downregulated genes in blue. **H.** Violin plot showing *Rgs5* normalized gene expression values (unique molecular identifier, UMI) for the pericyte cluster in 3 and 18 months old hearts. P value was calculated using *bimod* test.

To assess the transcriptional effects of ageing in cardiac pericytes, we analyzed single-nucleus RNA sequencing from 3 and 18 month old mice^13^, the first stage in which we detected a reduction of the pericyte coverage. We identified a well-defined pericyte cluster (**Figure 1E**) characterized by the expression of classical pericyte markers like *Pdgfrb*, *Cspg4*, *Rgs5*, *Abcc9*, *Kcnj8,* and *Mcam* (**Figure 1F**). Analysis of differentially expressed genes (DEG) between 3 and 18 month old pericytes revealed 46 DEG, 29 up and 17 down-regulated genes in old heart derived pericytes (**Figure 1G**). Gene ontology (GO) analysis of DEG showed a downregulation of genes related to electron transport chain, ATP metabolic process, and focal adhesion and downregulation of ERK1 and ERK2 cascade (**Supplementary** Figure 2A) suggesting that ageing impairs pericyte metabolism and pericyte ability to attach to the basement membrane. Furthermore, this analysis revealed that *Rgs5* was downregulated in the 18 month old pericytes (**Figure 1H**). This observation was validated by immunohistochemistry (**Supplementary** Figure 2B**, C**) and caught our attention since *Rgs5* is a very well-established pericyte marker gene^31^ (**Supplementary** Figure 2D). *RGS5* expression is also downregulated in primary human pericytes after prolonged cell culture, which induces replicative senescence and mimics the effect of ageing in vitro^32,33^ (**Supplementary** Figure 2E**, F**). The characterization of pericytes in vitro after prolonged culture revealed that size of the mural cells is increased (**Supplementary** Figure 2G**, H**). Similar the findings generated by the single-nucleus RNA sequencing from aged pericytes in vivo, RGS5 deficient pericytes showed a reduction of the focal adhesions (**Supplementary** Figure 2G**, I**). Although RGS5 has been studied in the heart in the context of pressure overload^34^, its role in cardiac pericytes at baseline and ageing conditions has not been studied.

### *RGS5* silencing in human pericytes reduces proliferation, migration and induces a change in the cell morphology

In order to gain insight into the role of RGS5 in pericytes, we depleted *RGS5* in cultured human pericytes from brain and placenta using siRNA (**Supplementary** Figure 3A**, B**). *RGS5* knockdown reduced pericyte cell cycle progression in vitro, as shown by a significant reduction of cells in S phase (**Figure 2A-D**). Furthermore, the knockdown of *RGS5* reduced pericyte migration (**Figure 2E-G**) and affected pericyte metabolism. In line with the gene ontology analysis of aged pericytes, metabolomics analysis of *RGS5*-deficient pericytes revealed reduced levels of tricarboxylic acid (TCA) cycle metabolites citrate, isocitrate, α-ketoglutarate, succinate and malate (**Supplementary** Figure 3C-G). Reduced TCA cycle activity was also linked to a decrease in cellular levels of NAD and NADP (**Supplementary** Figure 3H**, I**), as well as total ATP (**Supplementary** Figure 3J). These findings suggest that RGS5 deletion impairs pericyte mitochondrial metabolism. Pericytes are intimately associated with endothelial cells embedded within the vascular basement membrane^35^. To study the effects of *RGS5* knockdown on this interaction, we co-cultured both cell types and allowed endothelial cells and pericytes to form a vascular network in matrigel. Interestingly, upon *RGS5* knockdown, we detected a significantly increased number of morphologically altered pericytes that failed to align with the endothelial cell tubes (**Figure 2H-J**). This was further confirmed by analysis of pericyte morphology after *RGS5* knockdown. *RGS5* deficient pericytes, similar to pericytes after long term culture, became enlarged and presented a reduced number of focal adhesions (**Figure 2K-O**). Taken together, these observations indicate that RGS5 is a crucial regulator of pericyte cellular function and morphology in vitro.

**Figure 2.**
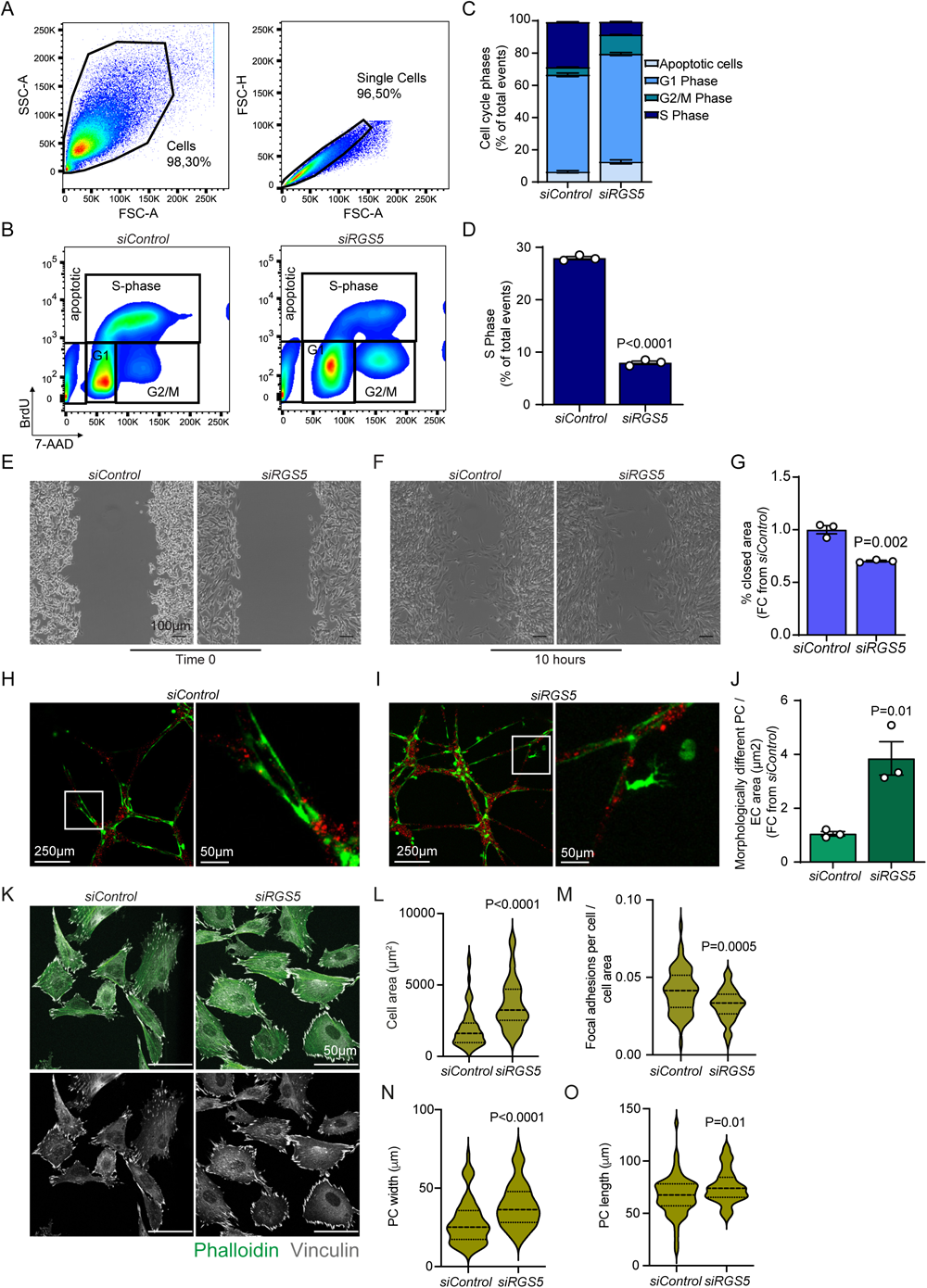
*RGS5* knockdown induces functional and morphological alterations in human pericytes. **A.** Gating strategy for BrdU proliferation assay using *siControl* treated hPC-PL. **B.** Representative FACS plots showing a reduction in the proliferation rate in *RGS5* knockdown pericytes (right) compared to *siControl* (left). **C.** Cell cycle phases distribution differences in hPC-PL between *siControl* and *siRGS5* conditions (*siControl:* apoptotic cells=7,76%, G1=57,90%, G2/M=4,85%, S=28,50%; *siRGS5*: apoptotic cells=12,10%, G1=67,40%, G2/M=12,20%, S=7,38%). **D.** Quantification of s-phase events (BrdU^+^ and 7AAD^+^) shows a significant reduction upon *RGS5* knockdown represented in Figure 2B. **E, F.** Representative images of hPC-PL migration at 0 and 10 hours timepoints comparing *siControl* and *siRGS5* conditions. **G.** Pericyte migration is reduced upon *RGS5* knockdown in hPC-PL. Values are expressed as percentage of closed area from time 0**. H, I.** Representative image of matrigel co-culture assay of pericytes (green) and endothelial cells (red). Morphologically different pericytes were found in *RGS5* knockdown sample. **J.** Quantification of morphologically different pericytes normalized to the total EC area (n=3). **K.** Immunofluorescence images of hPCPL showing the cytoskeleton (green) and the focal adhesions (white) indicating an increase in PC cell size upon *RGS5* knockdown. **L.** Quantification of the cell area. **M.** Quantification of the number of focal adhesion points per cell stained with vinculin and normalized to the cell area. **N.** Quantification of the pericytes cell width. **O.** Quantification of the pericytes cell length. **D, G, J, L-O.** Data are represented as mean ± S.E.M and P values are calculated using two tailed unpaired *t*-test.

### Pericyte-specific *Rgs5* deletion changes cardiac pericyte morphology in vivo

Although RGS5 has traditionally been used as a pericyte marker gene, its role in cardiac pericytes has not been studied. To do so, we bred mice bearing a conditional loss-of-function allele of *Rgs5*^9^ with *Pdgfrb-CreERT2* mice that express tamoxifen-inducible Cre recombinase in pericytes^12^. Cre expression was induced in adult animals (10 weeks old) and the hearts were subsequently analyzed 4 weeks and 8 weeks after tamoxifen injection (**Figure 3A**). Tamoxifen injection successfully deleted *Rgs5*-exon 3 in *Rgs5^fl/fl^*;*Pdgfrb-CreERT2 (Rgs5^ΔPC^*) mice (**Supplementary** Figure 4A) and strongly diminished the expression of RGS5 in the heart both at the gene expression and protein level (**Supplementary** Figure 4B-D). Similar to what we had observed in vitro, immunohistochemistry analysis of pericytes in controls and *Rgs5^ΔPC^* hearts revealed morphological alterations in these mural cells (**Figure 3B-F** and **Supplementary** Figure 4E-H). While control pericytes appeared organized with elongated thin branches following cardiac capillaries, *Rgs5^ΔPC^* pericytes appeared disorganized and presented wider cellular branches (**Figure 3B**, arrowheads). Surprisingly, we observed a significant increase of the total pericyte area in *Rgs5^ΔPC^* hearts, both 4 weeks and 8 weeks after tamoxifen injection (**Figure 3C**), accompanied by an increase in pericyte coverage only 4 weeks after *Rgs5* deletion (**Figure 3D**). Nevertheless, the total number of pericytes is not affected in the *Rgs5^ΔPC^* hearts (**Figure 3E**). We did not detect a different number of pericyte branches arising from the pericyte-cell body in *Rgs5^ΔPC^* hearts (**Figure 3F**), confirming that the increased pericyte area is due to a morphological change in pericytes characterized by wider pericyte branches rather than increased number of cells or branches. This observation is supported by our in vitro data in which we observed an increase in cellular area upon RGS5 knockdown.

**Figure 3.**
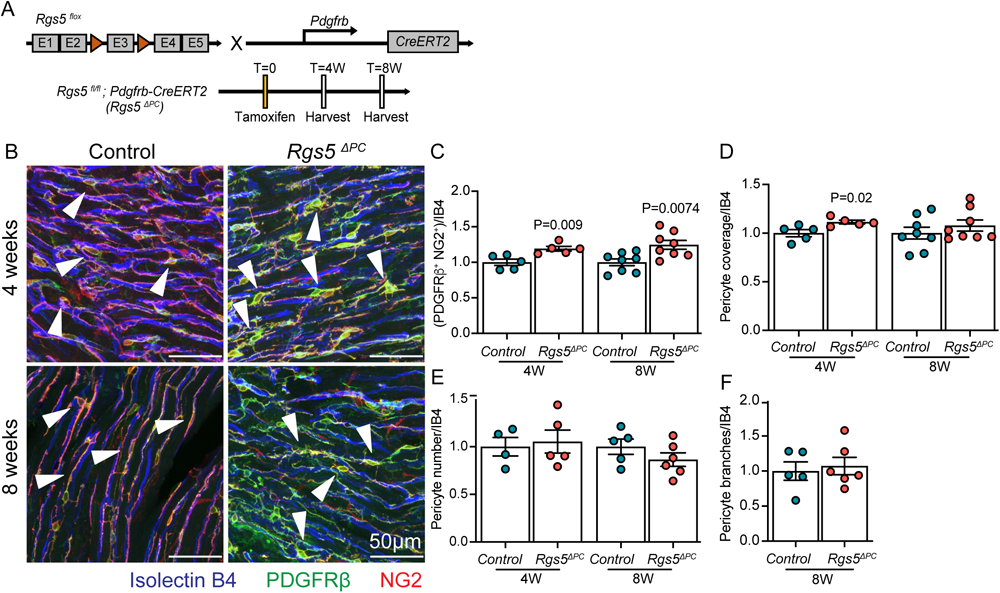
Pericyte total area is increased in *Rgs5^ΔPC^*hearts. **A.** Scheme of the experimental design. **B.** Immunofluorescence staining of left ventricular cross sections of *Control* and *Rgs5^ΔPC^* hearts 4 and 8 weeks after tamoxifen injection. Pericytes are indicated as double PDGFRβ and NG2 positive cells (arrowheads). **C.** Measurement of pericyte total volume shows significant increase in *Rgs5^ΔPC^* hearts both 4 and 8 weeks after tamoxifen injection. **D.** Measurement of pericyte coverage of the vasculature shows an increased pericyte volume in *Rgs5^ΔPC^* hearts 4 weeks after tamoxifen, back to normal 8 weeks after tamoxifen injection (Control and *Rgs5^ΔPC^* 8W, p=0.44). **E.** Pericyte cell bodies count normalized to total endothelial cell volume shows no difference between Control and *Rgs5^ΔPC^* (W4, n=4 and n=5 respectively, p=0.75; W8, n=5 and n=6 respectively, p=0.24). **F.** Pericyte branches count shows no change between Control and *Rgs5^ΔPC^* 8 weeks after tamoxifen injection (n=5 and n=6 respectively, p=0.70). **C-F.** All quantifications are normalized to the vasculature area and expressed as FC from Control. **C, D.** P values are calculated using two tailed unpaired *t*-test (Control vs *Rgs5^ΔPC^* at W4, n=5; Control and *Rgs5^ΔPC^* at W8, n=8) and represented as mean ± S.E.M. **E, F.** P values are calculated using two tailed unpaired *t*-test (Control vs *Rgs5^ΔPC^* at W4, n=4 and n=5 respectively; Control and *Rgs5^ΔPC^* at W8, n=5 and n=6 respectively) and represented as mean ± S.E.M.

### Loss of *Rgs5* in cardiac pericytes induces systolic dysfunction and increases collagen deposition in the heart

To characterize whether *Rgs5* deletion in pericytes would recapitulate some features of cardiac ageing, we studied cardiac structure and function in the *Rgs5^ΔPC^* mice. Echocardiography analysis of *Rgs5^ΔPC^* mice showed reduced left ventricular ejection fraction (EF) 4 weeks after tamoxifen injection, which was further reduced at 8 weeks (**Figure 4A, B**). Diastolic dysfunction as measured by the E/E’ ratio was increased by 20% but was not significantly different between control and *Rgs5^ΔPC^* mice (**Figure 4C**). These findings indicate that RGS5 deletion induces systolic but not diastolic dysfunction. Furthermore, impaired EF was accompanied by a significant increase in the thickness of the ventricular septum (**Figure 4D**), total left-ventricular (LV) mass (**Figure 4E, F**), and a significant reduction of diastolic LV volume (**Figure 4G**). These observations suggest that *Rgs5^ΔPC^* hearts become hypertrophic upon tamoxifen injection. Moreover, histological analysis revealed a significant increase in collagen deposition 8 weeks after tamoxifen injection in the myocardium (**Figure 4H, I**) but not in the perivascular region around bigger vessels (**Figure 4J, K**), indicative of an increased diffuse fibrosis in the *Rgs5^ΔPC^* hearts. In order to study whether this is a cardiac-specific phenotype, we analysed fibrosis in other pericyte-rich organs. Sirius red analysis revealed no increased collagen deposition in brain, lung, or kidney (**Supplementary** Figure 5A-D), indicating that the deletion of Rgs5 in pericytes induces fibrosis, specifically in the heart. However, the deletion of *Rgs5* in pericytes did not affect cardiomyocyte relative area (**Figure 4L, M**), vascular coverage (**Figure 5A, B**), capillary perimeter (**Figure 5A, C**) or vessel integrity analysed by the quantification of extravasated serum albumin or cadaverin injected in the tail vein (**Figure 5D-E** and **Supplementary** Figure 6A-C). We also did not detect hypoxic regions in the *Rgs5^ΔPC^* hearts (**Supplementary** Figure 6D). In order to determine whether the deletion of *Rgs5* would induce cellular senescence in the heart, we performed β-Galactosidase staining but we did not detect any positive area in the heart (**Supplementary** Figure 6E). Finally, the deletion of Rgs5 did not increase the number of infiltrating immune cells in the myocardium (**Figure 5F-I**). These data show that the deletion of RGS5 in pericytes affects pericyte morphology and impairs cardiac function and increases LV mass. In particular, RGS5 deletion induced collagen deposition indicating interstitial fibrosis in the myocardium.

**Figure 4.**
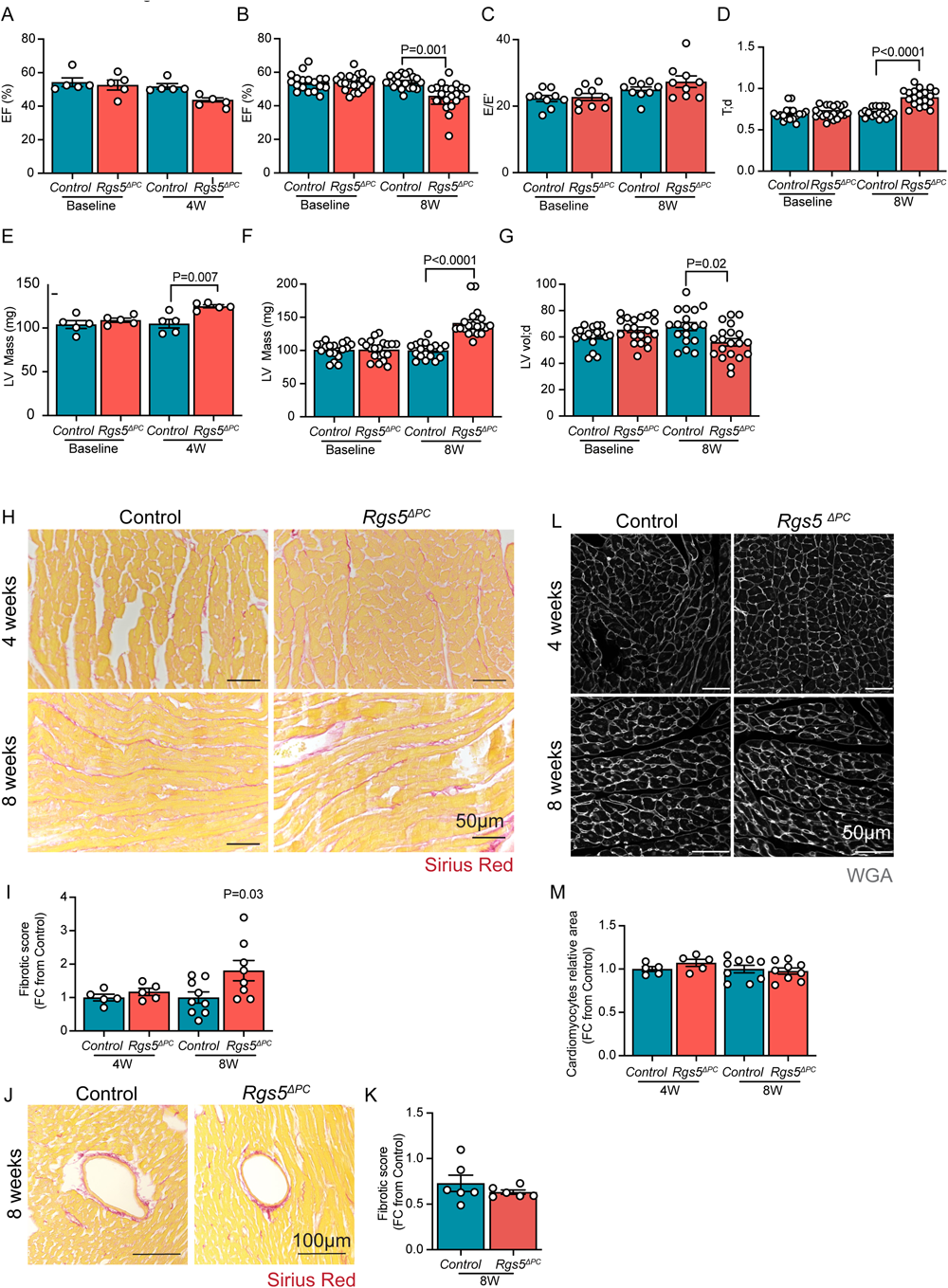
*Rgs5* deletion in pericytes compromises heart function and increases collaged deposition in the heart. **A-G.** Echocardiography analysis of control and *Rgs5^ΔPC^* mice. **A.** LV-EF in control and *Rgs5^ΔPC^* mice 4 weeks after tamoxifen injection (n=5; baseline control vs baseline *Rgs5^ΔPC^* P>0.99; baseline control vs control 4W P>0.99; baseline *Rgs5^ΔPC^* vs *Rgs5^ΔPC^* 4W P=0.08; control 4W vs *Rgs5^ΔPC^* 4W P=0.25). **B.** Echocardiography analysis of left ventricular ejection fraction in control and *Rgs5^ΔPC^* mice 8 weeks after tamoxifen injection (n=18 control, n=20 *Rgs5^ΔPC^*; baseline control vs baseline *Rgs5^ΔPC^* P>0.99; baseline *Rgs5^ΔPC^* vs *Rgs5^ΔPC^* 8W P=0.003; baseline control vs control 8W P>0.99). **C.** Echocardiography analysis of E/E’ value in control and *Rgs5^ΔPC^* mice 8 weeks after tamoxifen injection (n=9; baseline control vs baseline *Rgs5^ΔPC^* P=0.99; baseline control vs control 8W P=0.42; baseline control vs *Rgs5^ΔPC^* 8W, P=0.03). **D.** Echocardiography analysis of T;d value in control and *Rgs5^ΔPC^* mice 8 weeks after tamoxifen injection (n=18 control, n=20 *Rgs5^ΔPC^*; baseline control vs baseline *Rgs5^ΔPC^* P>0.99; baseline *Rgs5^ΔPC^* vs *Rgs5^ΔPC^* 8W P<0.001; baseline control vs control 8W P>0.99). **E.** Left ventricular mass measurements expressed in mg in control and *Rgs5^ΔPC^* mice 4 weeks after tamoxifen injection (n=5; baseline control vs baseline *Rgs5^ΔPC^* P=0.77; baseline *Rgs5^ΔPC^* vs *Rgs5^ΔPC^* 4W P=0.03 ;baseline control vs control 4W P=0.1). **F.** Left ventricular mass measurements expressed in mg in control and *Rgs5^ΔPC^* mice 8 weeks after tamoxifen injection (n=18 control, n=20 *Rgs5^ΔPC^*; baseline *Rgs5^ΔPC^* vs *Rgs5^ΔPC^* 8W P<0.001;baseline control vs baseline *Rgs5^ΔPC^* P>0.99; baseline control vs control 8W P>0.99). **G.** Echocardiography analysis of LV vol;d value in control and *Rgs5^ΔPC^* mice 8 weeks after tamoxifen injection (n=18 control, n=20 *Rgs5^ΔPC^*; baseline control vs baseline *Rgs5^ΔPC^* P>0.99; baseline control vs control 8W P=0.43; baseline *Rgs5^ΔPC^* vs *Rgs5^ΔPC^* 8W P=0.1). **A-G.** Data are represented as mean ± S.E.M and P values are calculated one-way ANOVA test. **H.** Representative pictures of Sirius Red collagen (red) staining of Control and *Rgs5^ΔPC^* left ventricular free wall of 4μm paraffin sections. **I.** Measurement of collagen area normalized to total area of the tissue. Fibrotic score of Control vs *Rgs5^ΔPC^* at 4W, p=0.28. **J.** Representative pictures of perivascular Sirius Red collagen (red) staining of Control and *Rgs5^ΔPC^* of 4μm paraffin sections. **K.** Measurement of perivascular collagen area normalized to the vascular lumen. Fibrotic score of Control vs *Rgs5^ΔPC^* at 4W, p=0.32.). **I, K.** Data are represented as mean ± S.E.M and P values are calculated two tailed unpaired *t*-test (Control vs *Rgs5^ΔPC^* at W4; Control and *Rgs5^ΔPC^* at W8). **L.** Wheat Germ Agglutinin (WGA) staining on 4μm paraffin sections of the left ventricular wall in Control and *Rgs5^ΔPC^* hearts 4 (n=5) and 8 weeks (n=9). **M.** Quantification of cardiomyocyte relative area shows no difference between Control and *Rgs5^ΔPC^* 4W (n=5, p=0.19), Control and *Rgs5^ΔPC^* W8 (n=9, n=8 respectively, p=0.69). **M.** Data are represented as mean ± S.E.M and P values are calculated using two tailed unpaired *t*-test (Control vs *Rgs5^ΔPC^* at W4; Control and *Rgs5^ΔPC^* at W8).

**Figure 5.**
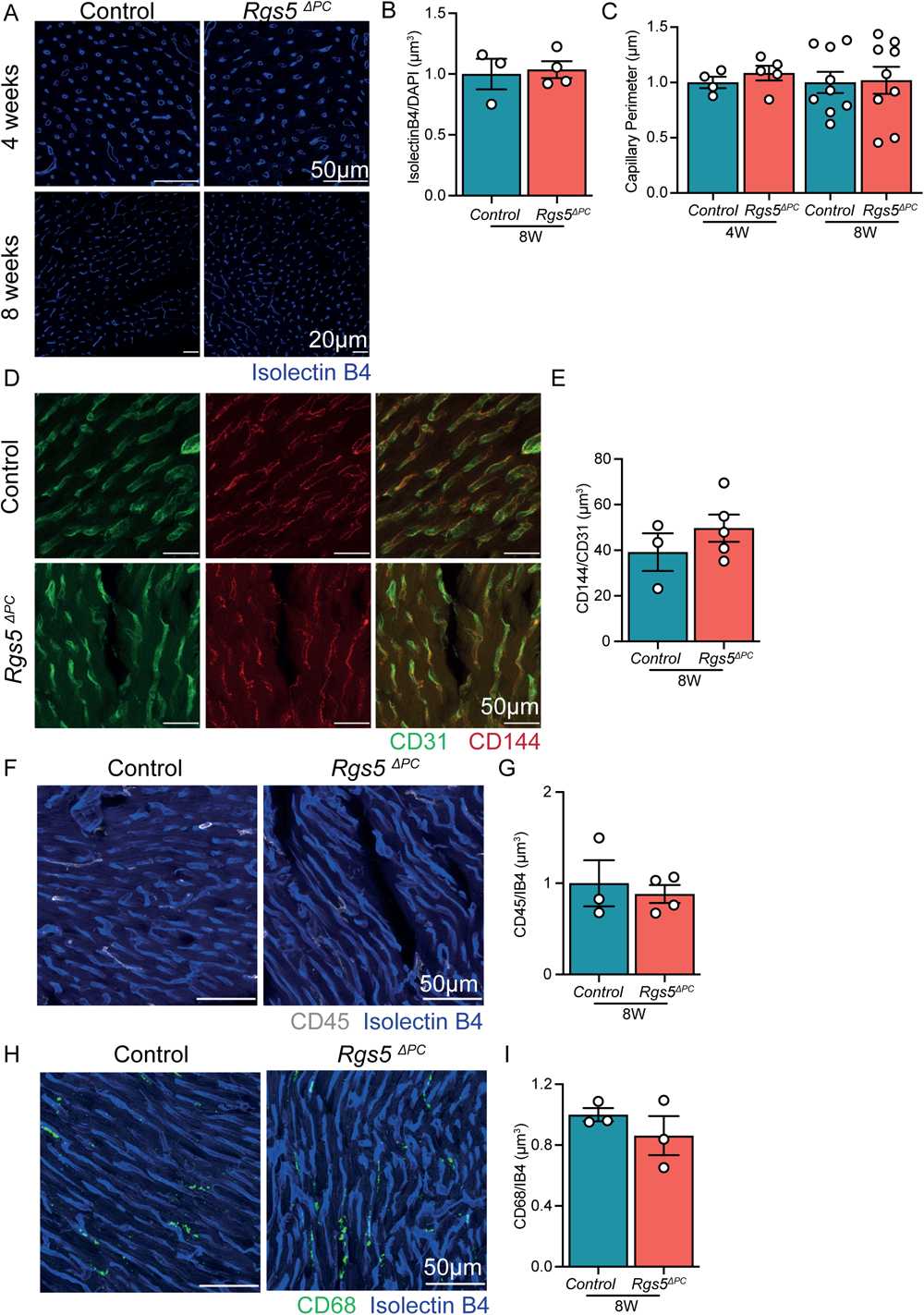
*Rgs5* loss in pericytes does not affect microvascular integrity. **A.** IsolectinB4 fluorescence staining of 4 µm paraffin sections of the free wall of the left ventricle in control and *Rgs5^ΔPC^* hearts 4 and 8 weeks after tamoxifen injection. **B.** Vascular coverage is not changed between control and *Rgs5^ΔPC^* hearts 8 weeks after tamoxifen injection (n=3 and n=4 respectively, P=0.80). **C.** Quantification of capillary perimeter shows no difference between control vs *Rgs5^ΔPC^* 4W (n=4, n=5 respectively, P=0.36), control and *Rgs5^ΔPC^* W8 (n=9 for both groups, P=0.9). **D, E.** Immunofluorescence staining and quantification of CD144 coverage (red) show no change in endothelial cell junctions between control and *Rgs5^ΔPC^* hearts 8 weeks after tamoxifen injection (n=3 and n=5 respectively, P=0.33). **F, G.** Immunofluorescence staining and quantification of CD45^+^ cells (white) show no change between control and *Rgs5^ΔPC^* hearts 8 weeks after tamoxifen injection (n=3 and n=4 respectively, P=0.65). **H, I.** Immunofluorescence staining and quantification of CD68^+^ cells (green) show no difference between control and *Rgs5^ΔPC^* hearts 8 weeks after tamoxifen injection (n=3, P=0.37). **E, G, I.** Data are shown as mean ± S.E.M. P value is calculated using two tailed unpaired *t*-test.

### RGS5 deficient pericytes show a pro-fibrotic gene expression profile and induce fibroblast activation

Fibroblasts are the main cell type responsible for the deposition of extracellular matrix in the heart^36,37^. Increased fibrosis can be mediated by an increase in fibroblast number or activation. Pericytes also secrete ECM and have been described to be a source of myofibroblasts following kidney injury^38^, in subretinal fibrosis^39^, or in cancer^40^. To study whether the deletion of *Rgs5* in pericytes would induce fibroblast expansion in the heart, we analysed the area covered by fibroblasts by staining PDGFRα, which is a well established marker for fibroblasts and not present in pericytes^41^. However, immunohistochemistry analysis of the PDGFRα area in *Rgs5^ΔPC^* hearts revealed no change in the area covered by fibroblasts (**Supplementary** Figure 7A-C). This observation suggests that the increase in collagen deposition is not due to an increase in fibroblast numbers, but rather it may be due to an induction of fibroblast activation. Similar to what was observed in the aged hearts, the total PDGFRβ positive area was increased in the *Rgs5^ΔPC^* hearts (**Supplementary** Figure 7A**, B, D**). Although PDGFRβ has been historically used to identify pericytes, it is also expressed in fibroblasts and stimulation of PDGFRβ in fibroblasts is known to induce collagen deposition^41^. In the *Rgs5^ΔPC^* hearts, the increase of PDGFRβ was detected outside pericytes, as we could detect an augmentation of PDGFRβ expression in NG2-negative population (**Supplementary** Figure 7E). However, the increase in PDGFRβ was detected in PDGFRα-expressing fibroblast resulting in a significant increase of PDGFRα and PDGFRβ double positive areas in the *Rgs5^ΔPC^* mice (**Supplementary** Figure 7F). Fibroblast activation was further confirmed by single-nuclei-RNA-sequencing. After annotating all major cardiac celltypes (**Figure 6A, B**), we first confirmed the reduction of *Rgs5* expression in pericytes and that it is not expressed in fibroblasts (**Figure 6B, C**). Following, we analyzed the expression of genes in cardiac fibroblasts (**Figure 6D-G**). Gene ontology analysis of differentially expressed genes in *Rgs5^ΔPC^* fibroblasts revealed a downregulation of genes related to the regulation of stress fiber assembly and RAS protein signal transduction (**Figure 6E**) and an upregulation of genes involved in collagen fibril organization and extracellular matrix organization (**Figure 6F**). In particular, we observed an upregulation of *Col3a1*, *Col15a1*, *Col14a1*, *Col6a2*, *Col5a3*, *Col4a3*, *Col4a2, Adamts12* and *Pxdn* (**Figure 6G**). Furthermore, we could confirm the increased expression of *Pdgfrb* (**Figure 6D**). Since the activation of PDGFRβ in fibroblasts induces collagen deposition and fibrosis^42,43^, we hypothesize that the deletion of RGS5 in pericytes drives the activation of cardiac fibroblasts thereby augmenting fibrosis possibly in a paracrine manner.

**Figure 6.**
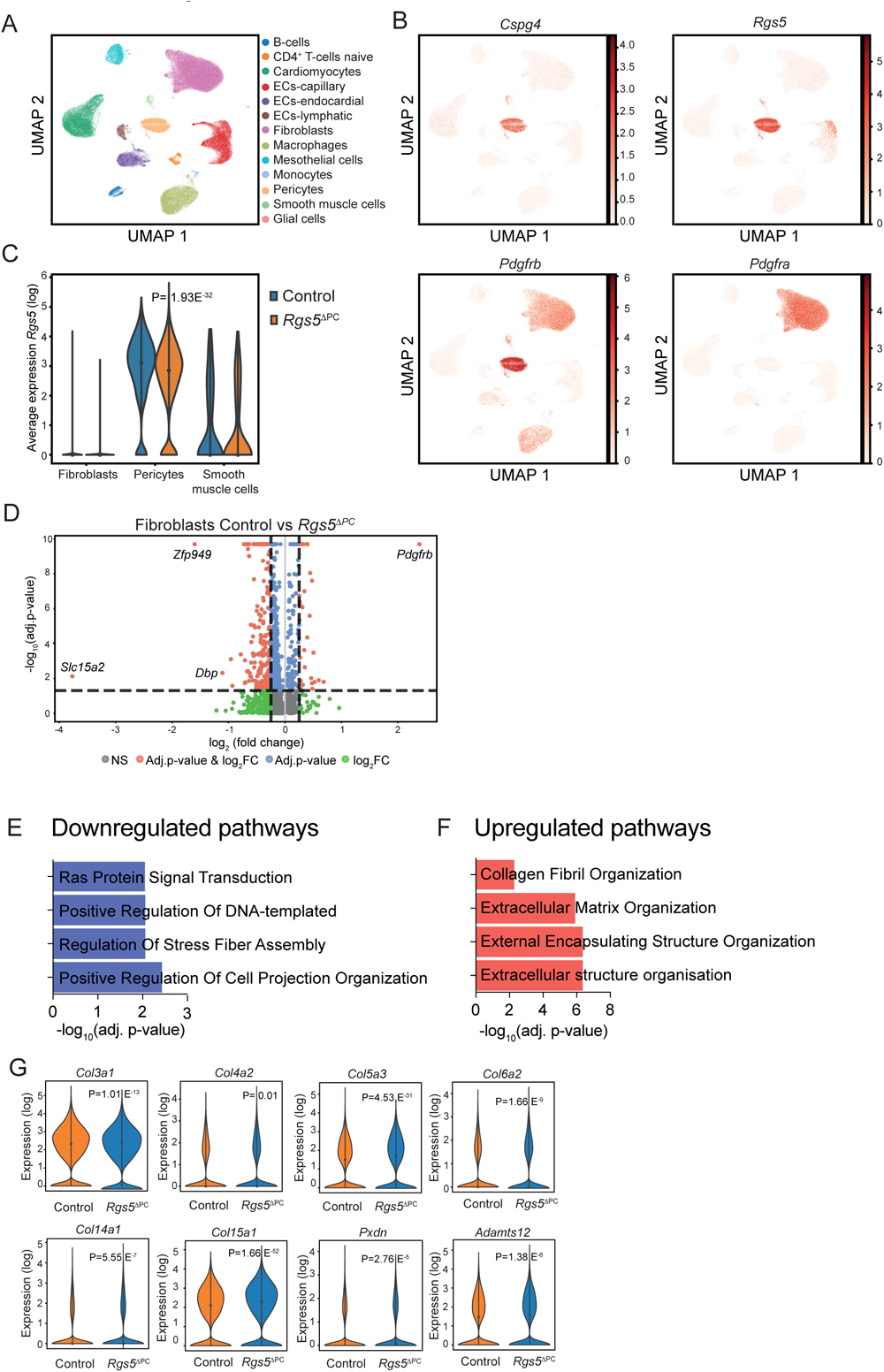
Single-nuclei sequencing confirmed fibroblast activation in *Rgs5^ΔPC^* hearts. **A.** Uniform Manifold Approximation and Projection (UMAP) plot showing cell-type specific clustering of all data points from cardiac single-nuclei sequencing. **B.** Feature plot showing gene expression of pericyte and fibroblast marker genes (*Pdgfrb, Cspg4, Rgs5, Pdgfra*). The colored scale bar indicates log-normalized gene expression level. **C.** Violin plot showing *Rgs5* normalized gene expression values (unique molecular identifier, UMI) for the pericyte, fibroblast and smooth muscle cells clusters. P value was calculated using Wilcoxon rank sum test. **D.** Volcano plot showing upregulated and downregulated genes in fibroblasts in Control and *Rgs5^ΔPC^* hearts. **E, F.** Gene Ontology (GO) enrichment analysis of significant downregulated and upregulated differentially expressed genes in fibroblasts in Control and *Rgs5^ΔPC^* hearts. Represented pathways are the top 4 in the list. **G.** Violin plot showing differentially expressed genes in fibroblasts between Control and *Rgs5^ΔPC^* hearts. P value was calculated using Wilcoxon rank sum test.

To understand the molecular mechanism by which RGS5 loss in cardiac pericytes may induce fibroblast activation, we performed bulk-RNA-sequencing in our *RGS5* deficient pericytes in vitro. Gene expression analysis revealed 1751 significant differentially expressed genes (DEG), 895 upregulated, 856 downregulated. GO analysis of regulated pathways showed an upregulation of genes involved in TGFβ regulation of extracellular matrix (ECM), ECM-receptor interaction and integrins cell surface interactions in angiogenesis (**Figure 7A**). On the contrary, the expression of genes related to cell cycle was downregulated (**Figure 7B**), confirming the reduction of cellular proliferation described above. Pro-fibrotic growth factors like *TGFB2* and *PDGFB*, which are known to induce fibroblast activation and fibrosis^44–47^, are upregulated upon *RGS5* knockdown (**Figure 7C**). *RGS5* silencing further induced structural ECM components like different collagens and laminins, genes related to ECM deposition or mesenchymal fate (e.g. *FN1*, *POSTN*, *SPARC, ACTA2)*. Moreover, pro-inflammatory receptors like *IL6R* and *IL7R* are upregulated upon *RGS5* knockdown (**Figure 7C**). RGS5 has been described as negative regulator of Gαq signalling ^9,48^. To investigate whether RGS5 regulates GPCR signalling in pericytes, we measured the Gαq-dependent inositol-1-phosphate (IP1) production upon RGS5 knockdown. Indeed, we observed a significant increase in IP1 production, suggesting that Gαq signalling is increased in pericytes upon RGS5 knockdown (**Figure 7D**). Furthermore, similar to the RGS5 knockdown, treating pericytes with the Gαq agonist U46619 induced the expression of TGFB2 and ACTA2 (**Figure 7E, F**). These data suggest that RGS5 knockdown unleash the inhibition of Gαq, which subsequently induces a pro-fibrotic pericyte phenotype. Finally, we tested whether the secretome of RGS5 deficient pericytes was sufficient to induce fibroblast activation. Therefore, we cultured human primary fibroblasts with the supernatant of control and RGS5 deficient pericytes. This experiment revealed that fibroblasts cultured with the conditioned medium of RGS5 deficient pericytes show an increased expression of α smooth muscle actin (αSMA) (**Figure 8A, B**), a sign of fibroblast activation ^49^. Together, our pericytes bulk RNA sequencing and Gαq experiments suggested that RGS5 may mediate the activation of fibroblasts via the increase expression of TGFβ2. To test this hypothesis, we cultured human primary fibroblasts with the supernatant of control and RGS5 deficient pericytes in the presence of TGFβ neutralizing antibodies. This treatment rescued the activation of fibroblasts cultured with conditioned medium from RGS5 deficient pericytes (**Figure 8C, D**). In conclusion, these data suggest that pericytes not only activate the Gαq-TGFβ axis, but indirectly contribute to the increased fibroblast activation and deposition of extracellular matrix. These two features may contribute together to the increased fibrotic score observed in the *Rgs5^ΔPC^* hearts.

**Figure 7.**
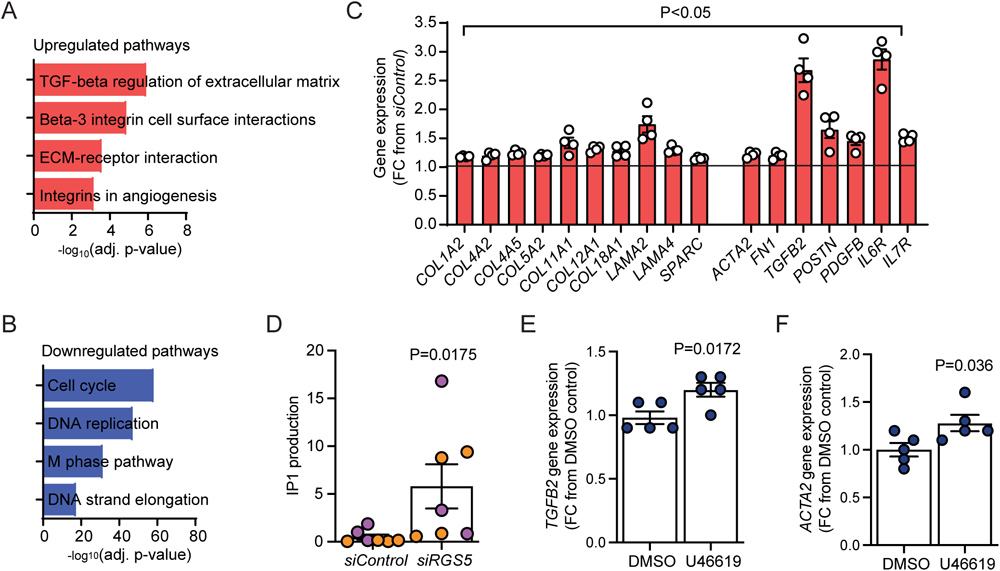
*RGS5* knockdown induces a pro-fibrotic phenotype in human pericytes. **A, B.** Gene Ontology (GO) enrichment analysis of significant differentially expressed genes after *RGS5* knockdown. Represented pathways are the top 4 in the list. n=4 replicates from different hPC-PL donors. **C.** List of significant upregulated genes upon *RGS5* knockdown involved in fibrosis, ECM deposition and inflammation**. D.** IP1 production in hPC-PL after RGS5 knockdown. Different colour represents two different hPC-PL lots. **E, F.** RT-qPCR analysis of *TGFB2* and *ACTA2* gene expression in hPC-PL after U46619 treatment (n=5). **D-F.** Values are shown as FC from siControl. **D, E, F.** Data are represented as mean ± S.E.M and P value is calculated using two tailed unpaired *t*-test.

**Figure 8.**
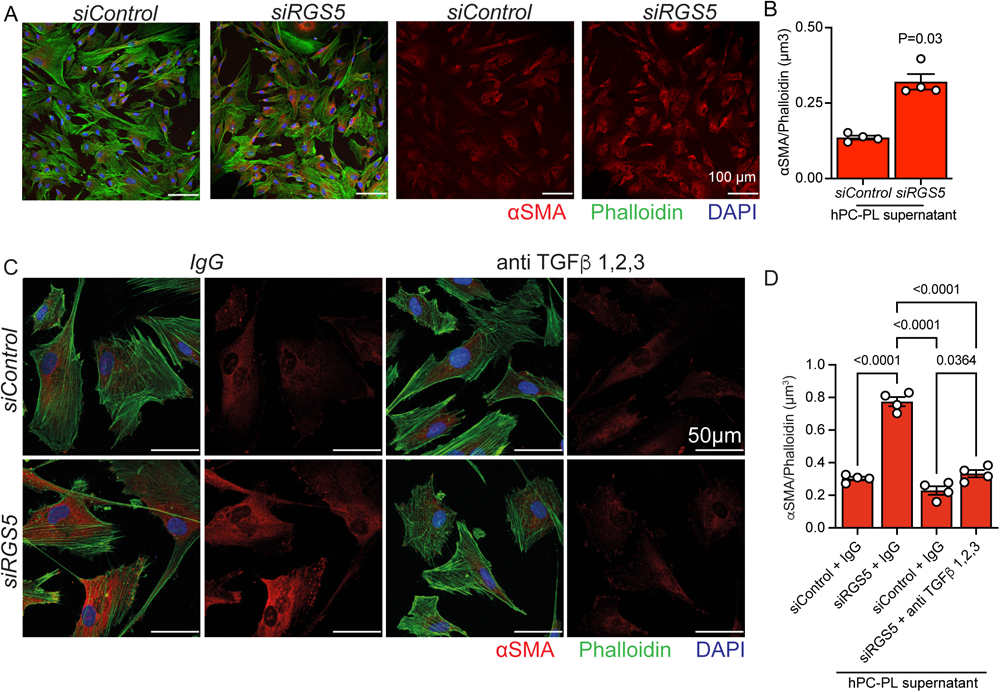
RGS5 deficient pericytes activate fibroblasts via TGFβ. **A.** Immunofluorescence staining of human cardiac fibroblasts showing an increase in αSMA^+^ activated fibroblasts treated with the supernatant from *siRGS5* and *siControl* hPC-PL. **B.** Quantification of αSMA^+^ volume normalized to phalloidin (n=4). Data are represented as mean ± SEM and P values are calculated using two tailed unpaired *t*-test. **C.** Immunofluorescence staining of HCF treated with RGS5 deficient PC supernatant in the presence of TGFβ 1,2,3 neutralising antibody. Neutralising antibody rescued HCF activation. **D.** Quantification of αSMA^+^ volume normalized to phalloidin cell area (n=4). (siControl + IgG vs siControl + TGFβ 1,2,3, P=0.17; siControl + IgG vs siRGS5 + TGFβ 1,2,3, P=0.79). Data are represented as mean ± S.E.M and P values are calculated using one-way ANOVA test.

## Discussion

We have studied the effect of ageing on cardiac pericytes and the role of RGS5 in these mural cells. We demonstrated that deleting RGS5 in pericytes induces a pro-fibrotic response, characterized by increased gene expression of different structural components of the extracellular matrix, but also of pro-fibrotic growth factors *TGFB2* and *PDGFB*. These two growth factors have already been described to play a role in the activation and induction of extracellular matrix deposition of fibroblasts^44–47^. The pro-fibrotic gene signature is confirmed in vitro and accompanied by a reduction of cellular proliferation, cellular migration, and morphological changes of pericytes. Furthermore, we have confirmed that the secretome of RGS5 deficient pericytes induces the activation of fibroblasts in a Gαq-TGFβ dependent mechanism.

In vivo, the deletion of *Rgs5* induces similar morphological changes to the pericytes, causes cardiac dysfunction, and leads to an increase of collagen deposition and fibrosis in the heart, which are hallmarks of ageing^50,51^. Interestingly, this phenotype was only detected in the heart suggesting that the role of RGS5 in regulating fibrosis is heart specific. The increased collagen deposition was accompanied by the activation of the fibroblast population in the heart as shown by the increased expression of PDGFRβ. *PDGFRB* is a direct transcriptional target of TGFβ^52^ and the activation of PDGFRβ signaling in fibroblasts can drive fibrosis^42,43^, suggesting that, similar to what we observed in vitro, the secretion of TGFβ2 by pericytes may be responsible for the activation of the cardiac fibroblasts. Moreover, we observed that RGS5 deficient pericytes increase *ACTA2*, *FN1*, *POSTN* and collagens gene expression and also cell area, a feature that in fibroblasts has been related to activation ^53^. These experiments suggest that upon RGS5 loss, pericytes become activated and participate actively in the fibrotic process.

Although we could not detect cardiomyocyte hypertrophy, the deletion of *Rgs5* induced the thickness of the ventricular septum, the increase of the LV mass in the heart, and the reduction of the LV diastolic volumes. We hypothesize that this is driven by the excessive accumulation of collagens as myocardial fibrosis has been shown to be related to increase LV mass^54^. Myocardial fibrosis can also contribute to sudden cardiac death, ventricular tachyarrhythmia, LV dysfunction, and in the end, heart failure^55^. In this sense, we have observed that the deletion of RGS5 in pericytes compromises LV ejection fraction, a major readout of LV function. Our data also indicate that pericytes themselves may participate in the increased extracellular matrix deposition observed in the mutant hearts.

The deletion of RGS5 did not induce a reduction of the pericyte area or coverage in vivo despite reducing cellular proliferation in vitro. This discrepancy might be explained by our experimental design in which we only followed *Rgs5^ΔPC^* up to eight weeks. The heart is a relative quiescent organ with little cellular proliferation in vivo^56–58^. We hypothesize that the *Rgs5^ΔPC^* mice, similar to the aged mice, would also show reduced pericytes numbers in the heart after prolonged deletion. Alternatively, RGS5 deficiency might be partially compensated in vivo.

Although RGS5 is also expressed in smooth muscle cells, no difference in blood systolic and diastolic pressure, and mean arterial pressure were observed in the global *Rgs5* KO^59^ and in smooth muscle cell specific *Rgs5* knock-out mice^9^ under baseline conditions. Furthermore, we detected difuse but not perivascular fibrosis in the *Rgs5^ΔPC^* hearts. Opposite to our results in pericytes, *RGS5* deficient smooth muscle cells show increased cellular proliferation^9^ indicating that RGS5 might play different roles depending of the cellular context and supporting our interpretation that the phenotype that we observed in the *Rgs5^ΔPC^* hearts is caused by the deletion of *Rgs5* in pericytes. In this sense, although *Pdgfrb* is expressed in fibroblasts and thus, the recombinase Cre would be expressed in these mesenchymal cells, we have shown that *Rgs5* is not expressed in fibroblasts further supporting our hypothesis that the observed phenotype is due to the deletion of RGS5 in pericytes.

In conclusion, our results identified the reduction of pericyte RGS5 as an indication of cardiac ageing and enlightens the importance of pericytes maintaining cardiac homeostasis. We have shown for the first time that RGS5 is crucial for the maintenance of pericyte function preventing the activation of a pro-fibrotic gene expression signature in the heart. The loss of RGS5 in pericytes drives an entropic state of these mural cells characterized by morphological changes, excessive ECM deposition, and secretion of pro-fibrotic growth factors that is pernicious for the heart.

### Limitations

Although our study offers valuable insights into the role of cardiac pericytes and Rgs5 in the process of cardiac ageing, our animal model does not completely mimic cardiac ageing. We induced a total deletion of Rgs5 in pericytes rather than a progressive and modest reduction observed in aged animals, and we did not observe any effect on diastolic function, an important hallmark of cardiac ageing. Furthermore, we only followed the animals for 8 weeks, which might have not sufficient to observe the long term chronic effect of Rgs5 deletion.

A second limitation is that we have not performed our in vitro experiments using cardiac pericytes. As it has been shown in this study, pericytes have different functions in different organs. Cardiac pericytes are difficult to isolate and culture and it is not clear, whether the organotypical differences that pericytes possess in vivo may be maintained in vitro once cultured. Nevertheless, our in vitro studies recapitulate the observations made in vivo. For this reason, it may be justified to use brain and placenta pericytes and HUVEC.

### Novelty and Significance

What is known

- Cardiac microvasculature dysfunction is associated with heart failure and cardiac disease.
- Pericytes are capillary-associated mesenchymal cells crucial for the maintenance of vascular homeostasis.
- RGS5 is an activator of GTPases known to regulate proliferation and contraction on smooth muscle cells.

What new information does this article contribute?

- Pericyte coverage is reduced in the ageing heart.
- Aged pericytes are characterized by a reduction of *Rgs5* gene expression.
- RGS5 loss of function induces a pro-fibrotic response in pericytes, fibrosis and systolic dysfunction in the heart.
- RGS5 loss induces fibroblast activation in a Gαq-TGFβ dependent mechanism.

Ageing reduces pericyte coverage in the heart and the reduction of pericyte RGS5 is an indication of pericyte ageing. The deletion of RGS5 in pericytes causes cardiac dysfunction, increases the left ventricular mass, and induces myocardial fibrosis, one of the hallmarks of cardiac ageing. Mechanistically, this work suggests that the reduction of RGS5 induces a pro-fibrotic gene expression signature characterized by the expression of different extracellular matrix components and TGFβ2 in a Gαq dependent mechanism that induce the activation of fibroblasts in a paracrine manner and fibrosis in the myocardium.

## Acknowledgements

This study was supported by the DFG SFB1531 project B5 to GL, project B4 to SD, project A4 to NW, and project S1 to IF. The authors would like to thank Katja Schmitz for her technical help with the echocardiography and Thomas Korff for the kind shipping of the *Rgs5^flox^* mice. A.T. would like to especially thank Veronica Larcher, Andreas W Heumüller and Mariana Shumliakivska for their support and time during ideas brainstorming.

## Autor contributions

A.T. performed in vitro and in vivo experiments and data analysis. L.S.T. and D.R.M. performed bioinformatic analysis. A.F. performed and analysed echocardiography data. M.M-R. and B.N.T performed histochemistry experiments. L.R.V. performed immunocitochemisty analysis. J.N. performed human vascular brain pericytes experiments. S.F.G. performed single-nuclei isolation of control and *Rgs5^ΔPC^* hearts. M.M. analysed FACS data. J.K. supported IP1 measument experiments. S.K. performed metabolomics experiment. B.S. performed single-nuclei library preparation. S.G. performed bulk RNA sequencing analysis. W.A. supported single-nucleus RNA sequencing experiments. D.J. supported bioinformatic analysis. I.F. supported metabolomics analysis. N.W. generated the *Rgs5^flox^* mouse line. A.T., G.L., S.D. conceived and designed experiments. A.T. and G.L. wrote the manuscript. All authors reviewed and commented on the manuscript.

---

**Supplementary Figure 1.**
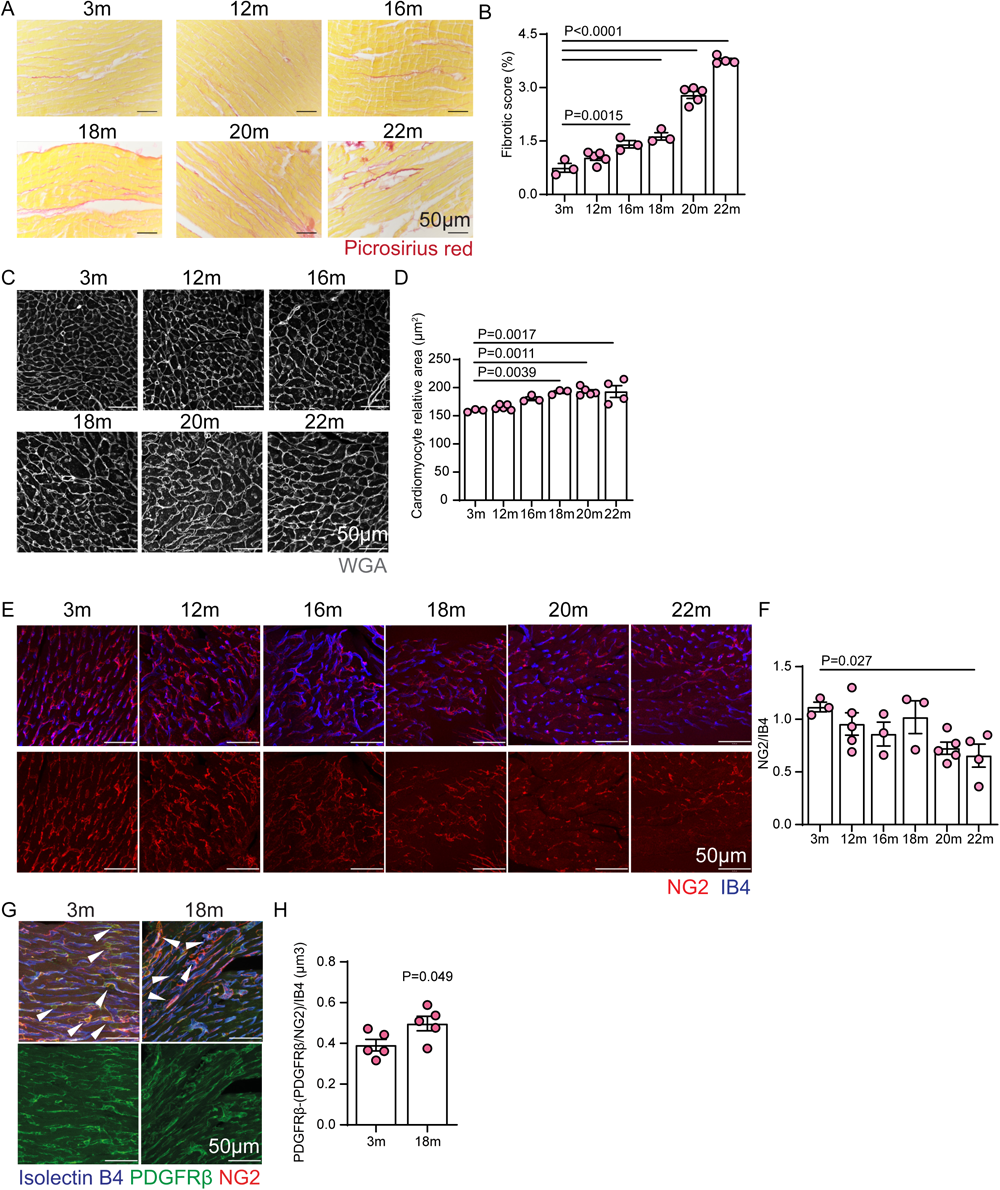
Cardiac ageing induces fibrosis and hypertrophy and reduces pericyte area. **A.** Representative pictures of Sirius Red (red) staining on 4μm paraffin sections of 3, 12, 16, 18, 20, 22 months C57BL/6 murine free wall of the left ventricle. **B.** Measurement of Sirius Red area normalized to total area of the tissue. Fibrotic score of 3m vs 12m (p=0.16). **C.** Wheat Germ Agglutinin (WGA) staining on 4μm paraffin sections of the free wall of the left ventricle in 12, 16, 18, 20, 22 months. **D.** Quantification of cardiomyocyte relative area shows no difference between 3m and 12m hearts (p=0.86) and between 3m and 16m hearts (p=0.08). **E.** Immunofluorescence staining of left ventricular cross sections of 3m, 20m and 22m hearts stained for NG2 and Isolectin B4. **F.** Quantification of NG2 normalized to the vasculature area. NG2 area 3m vs 12m (p=0.68), 3m vs 16m (p=0.38), 3m vs 18m (p=0.95), 3m vs 20m (p=0.055). **G.** Immunohistochemistry of left ventricular cross sections of young (3 months) and old (18 months) C57BL/6 murine hearts. Pericytes are identified as double PDGFRβ and NG2 positive cells (arrowheads). **H.** Quantification of PDGFRβ outside of the pericyte population normalized to the vasculature area, n=5 per condition. Data are shown as mean ± S.E.M. P value is calculated using two tailed unpaired *t*-test. **B, F.** Data are expressed as FC from Control. **B, D, F.** Data are represented as mean ± S.E.M and P values are calculated using ordinary one-way ANOVA test followed by Tukey’s multiple comparisons test. 3m, n=3, 12m, n=5, 16m, n=3, 18m, n=3, 20m, n=5, 22m, n=4.

**Supplementary Figure 2.**
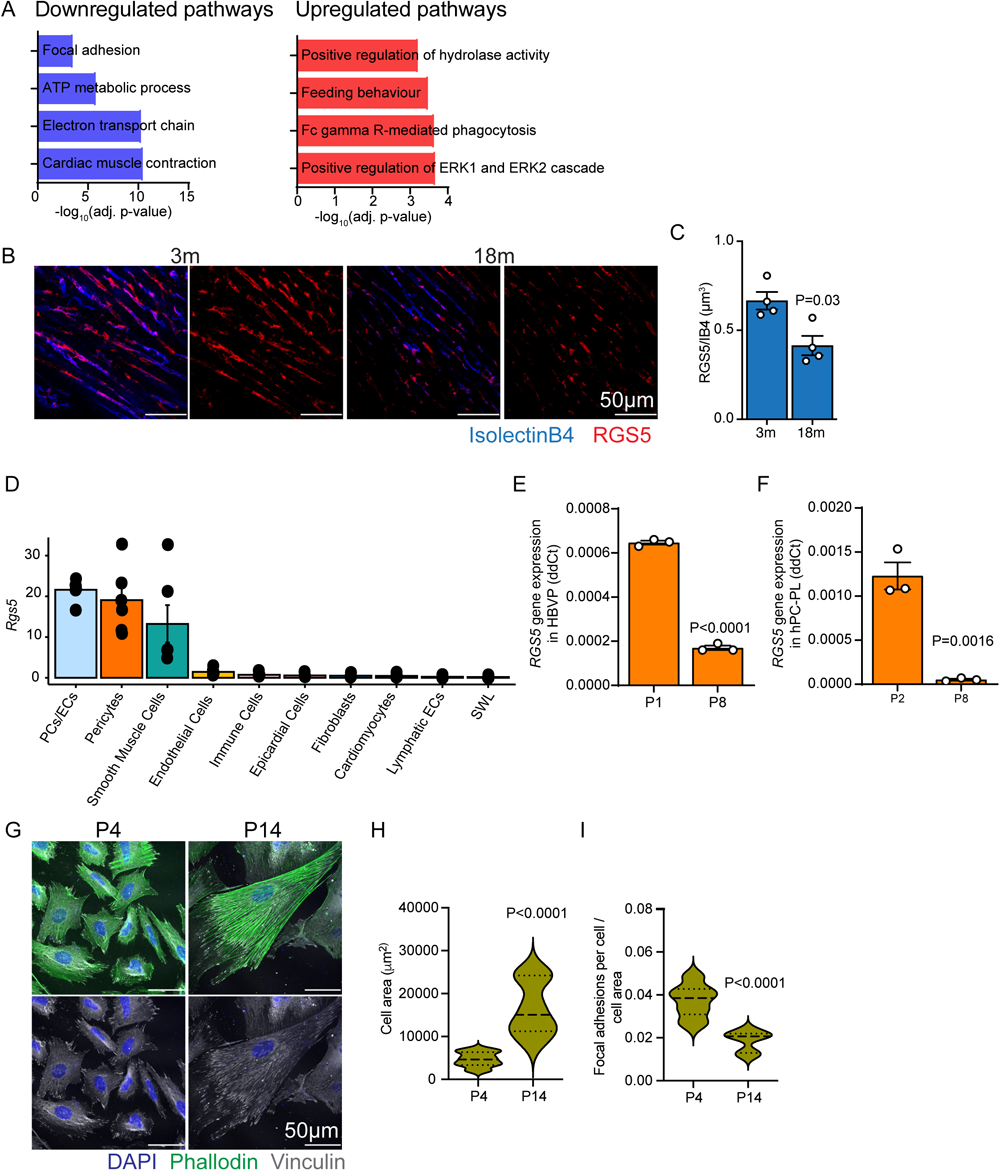
Cardiac ageing affects on pericyte gene expression. **A.** Gene Ontology (GO) enrichment analysis of significant differentially expressed gene between young and old pericytes. Represented pathways are the top 4 in the list. **B.** Immunohistochemistry of left ventricular cross sections of young (3 months) and old (18 months) C57BL/6 murine hearts showing a reduction in RGS5 in the old heart. **C**. Quantification of RGS5 normalized to the vasculature area, n=4 per condition. Data are shown as mean ± S.E.M. P value is calculated using two tailed unpaired *t*-test. **D.** *Rgs5* gene expression abundance in all cell types in the murine heart (n=6). **E, F.** RT-qPCR analysis of *RGS5* gene expression in early passage (P1, P2) and late passage pericytes (P8) in HBVP and hPC-PL (n=3). Data are represented as mean ± S.E.M and P values are calculated using two tailed unpaired *t*-test. **G.** Immunofluorescence images of early passage (P4) and late passage (P14) hPCPL showing the cytoskeleton (green) and the focal adhesions (white) indicating an increase in pericyte cell size upon *RGS5* knockdown. **H.** Quantification of the cell area. **I.** Quantification of the number of focal adhesion points per cell stained with vinculin and normalized to the cell area. **H, I.** Data are represented as mean ± S.E.M and P values are calculated using two tailed unpaired *t*-test.

**Suplemmentary Figure 3.**
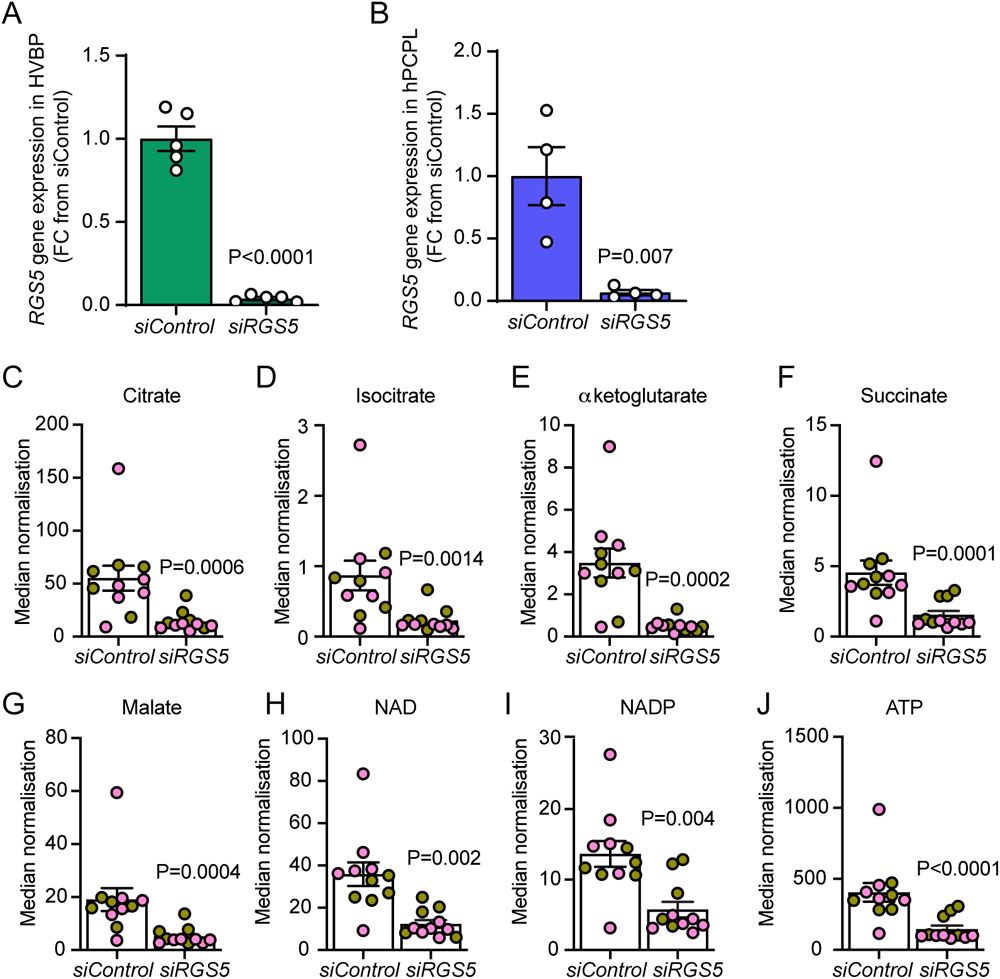
*RGS5* knockdown in human pericytes. **A, B.** RT-qPCR gene expression analysis of *RGS5* showing efficient *RGS5* silencing in human brain vascular pericytes (a, n=5) and human placenta pericytes (b, n=4). Data are represented as mean ± S.E.M and P values are calculated using two tailed unpaired *t*-test. **C-J**. Metabolomics analysis of RGS5 deficient pericytes. Metabolite concentration is normalized to total protein concentration. Data are represented as mean ± S.E.M and P values are calculated using two tailed unpaired *t*-test.

**Supplementary Figure 4.**
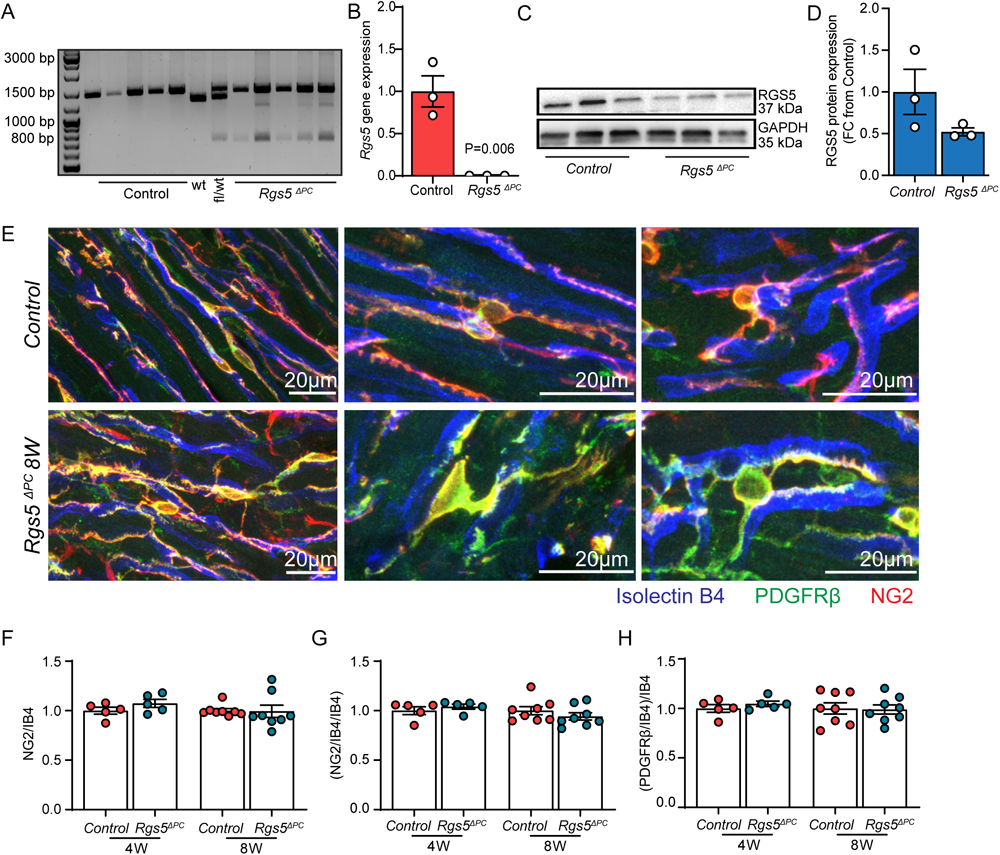
NG2 total area is unchanged in *Rgs5 ^ΔPC^* hearts. **A.** Electrophoresis analysis of PCR amplified Genomic DNA isolated from the apex of Control and *Rgs5^ΔPC^* hearts 4W after tamoxifen injection. Wildtype (wt) and heterozygote (fl/wt) controls were included in the analysis. **B.** RT-qPCR analysis of *Rgs5* gene expression in 8W Control and *Rgs5^ΔPC^* hearts. Data are represented as mean ± S.E.M and P values are calculated using two tailed unpaired *t*-test **C.** Western blot of protein lysates isolated from the apex of 8W Control and *Rgs5^ΔPC^* hearts. **D.** Quantification of RGS5 total protein in 8W Control and *Rgs5^ΔPC^* hearts (P=0.16). **E.** Representative high magnification immunofluorescence images of pericytes acquired in left ventricular cross sections of 8W Control and *Rgs5^ΔPC^* hearts. **F, G.** NG2 total volume (4W, p=0.21; 8W, p=0.16) and NG2 coverage (4W, p=0.30; 8W, p=0.31) of the vasculature are unchanged between Control and *Rgs5^ΔPC^* hearts. **H.** PDGFRβ coverage of the vasculature shows no difference between Control and *Rgs5^ΔPC^* hearts (4W, p=0.36, 8W, p=0.88). **F-H.** Data are normalized to the vasculature area and expressed as FC from Control. Data are represented as mean ± S.E.M and P values are calculated using two tailed unpaired *t*-test (Control vs *Rgs5^ΔPC^* at W4, n=5; Control and *Rgs5^ΔPC^* at W8, n=8).

**Supplementary Figure 5.**
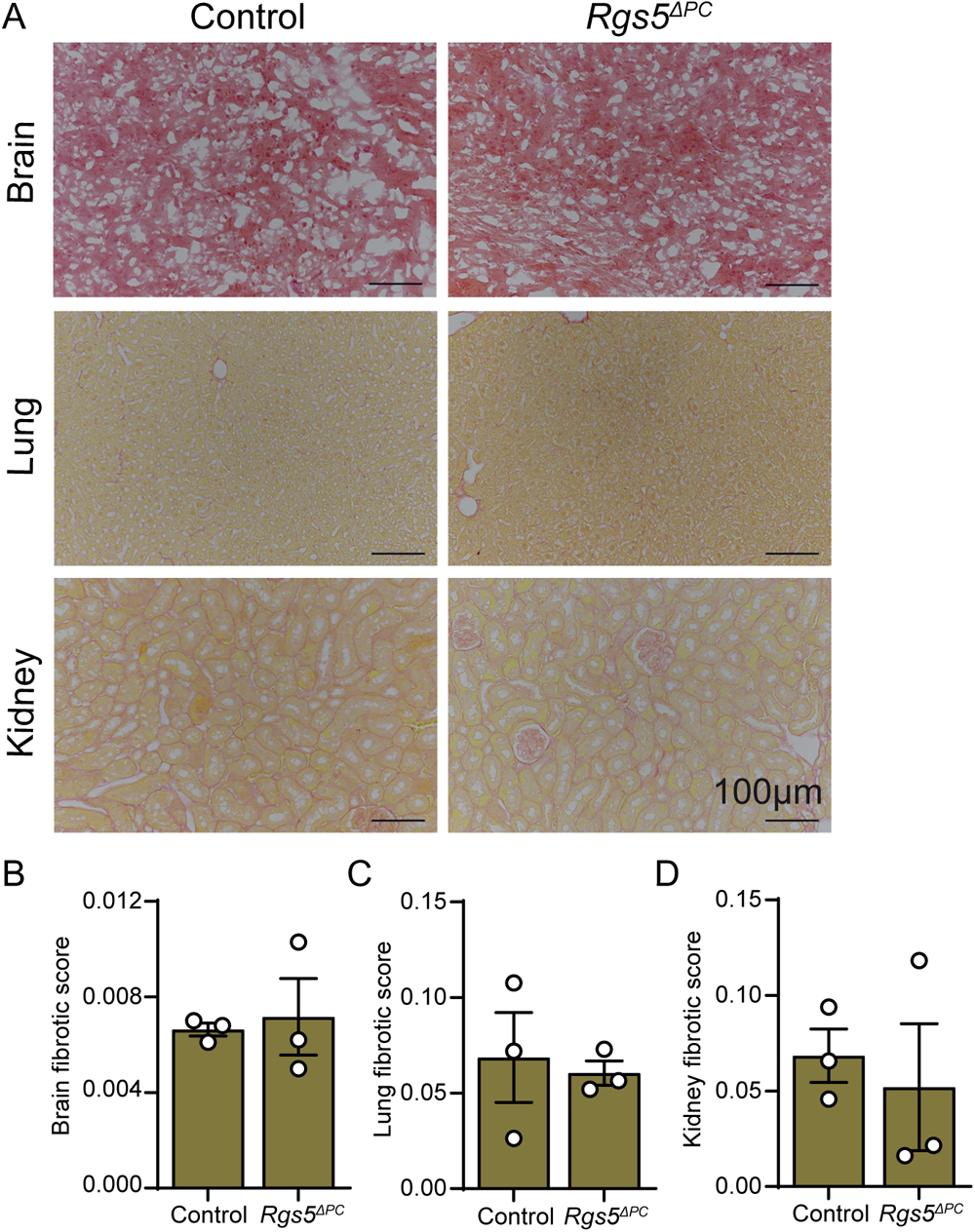
Loss of *Rgs5* in cardiac pericytes does not induce fibrosis in brain, lung and kidney. **A.** Representative pictures of Sirius Red collagen (red) staining of 4μm paraffin sections of Control and *Rgs5^ΔPC^* brain, lung and kidney. **B-D.** Measurement of collagen area normalized to total area of the tissue. Data are represented as mean ± S.E.M and P values are calculated using two tailed unpaired *t*-test (brain control vs *Rgs5^ΔPC^*, P=0.76; lung control vs *Rgs5^ΔPC^*, P=0.67; kidney control vs *Rgs5^ΔPC^*, P=0.76).

**Supplementary Figure 6.**
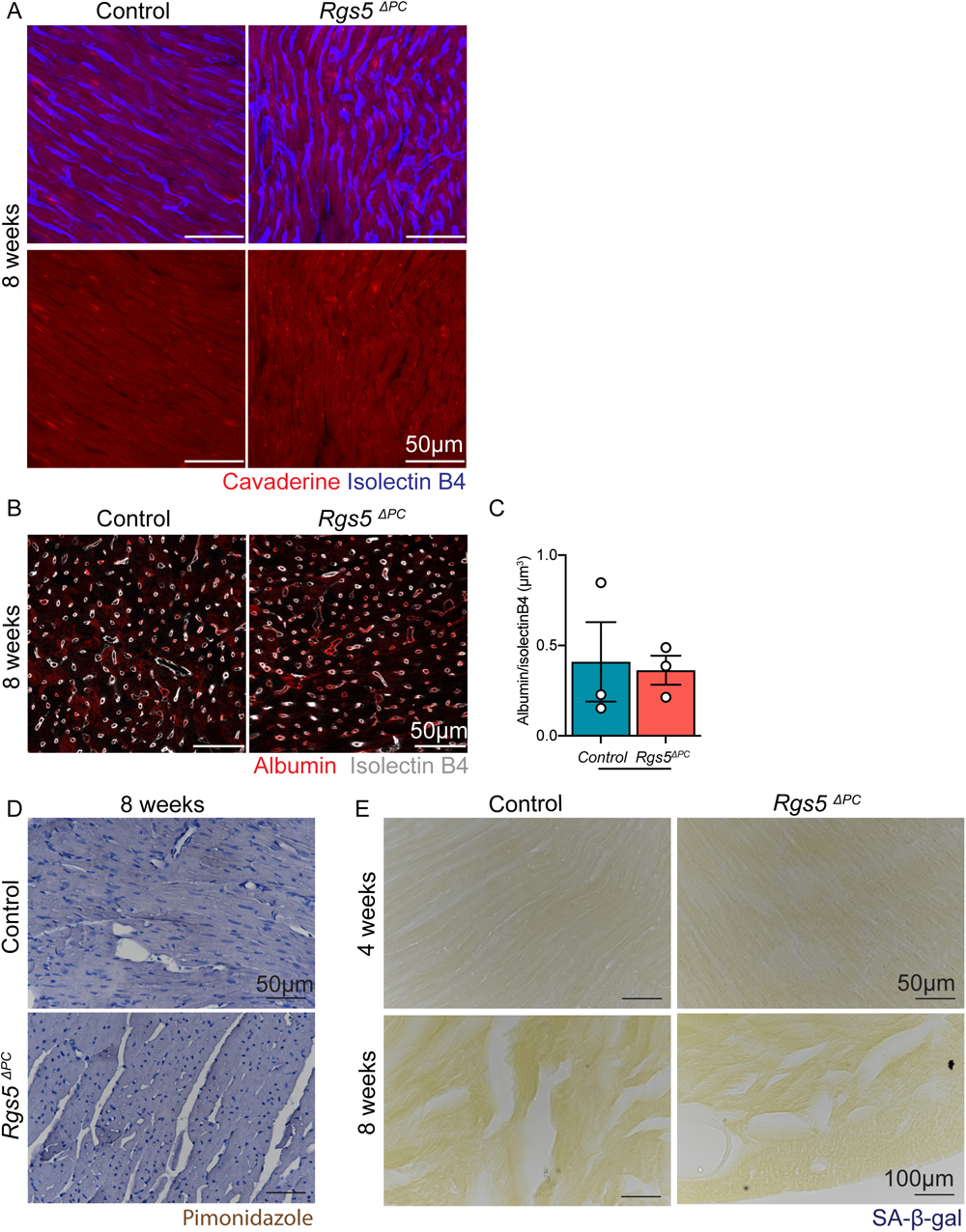
*Rgs5* deletion does not affect vascular leakage, senescence, or hypoxia in the heart. **A.** Analysis of cadaverine (red) extravasation in control and *Rgs5^ΔPC^* 8 week after tamoxifen injection shows no microvascular leakage. **B, C.** Immunofluorescence staining, and quantification of Albumin extravasation (red) shows no change between Control and *Rgs5^ΔPC^* 8 week after tamoxifen injection (n=3, p=0.85). Data are shown as mean ± SEM. P value are calculated using two tailed unpaired *t*-test. **D.** Hypoxia assay using Pimonidazole revealed no hypoxic area in the tissue of Control and *Rgs5^ΔPC^* 8 weeks after tamoxifen injection. **E.** β Galactosidase staining of Control and *Rgs5^ΔPC^* 4 and 8 weeks after tamoxifen injection shows no senescent cells in the heart. **H, I.** Immunofluorescence staining and quantification of CD45^+^ cells (white) show no change between Control and *Rgs5^ΔPC^* hearts 8 weeks after tamoxifen injection (n=3 and n=4 respectively, p=0.65).

**Supplementary Figure 7.**
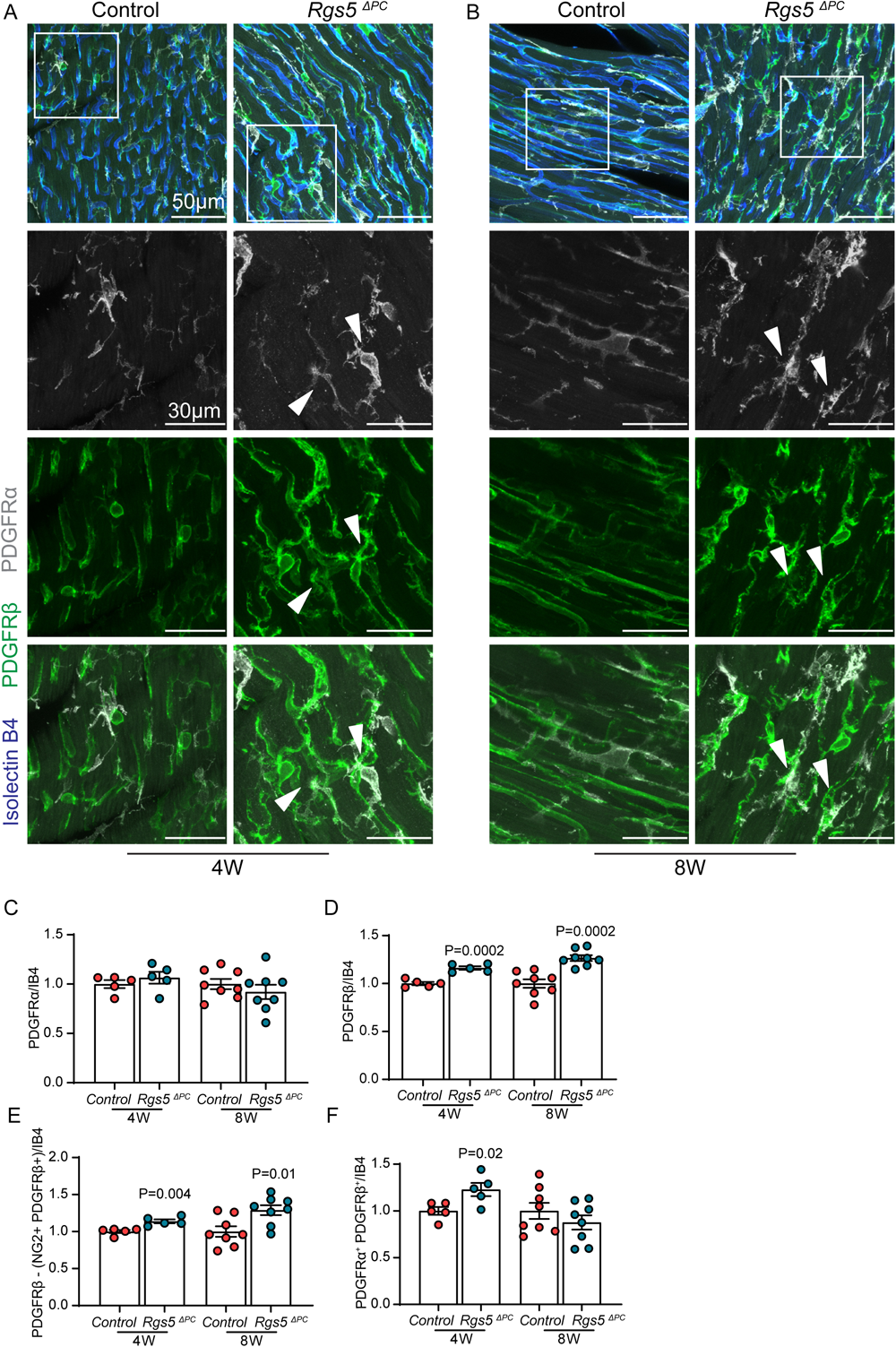
*Rgs5* deletion contributes to induce fibroblast activation in vivo. **A, B.** Immunostaining of left ventricular cross sections of 4W and 8W Control and *Rgs5^ΔPC^* hearts showing PDGFRα^+^ fibroblasts (white) and PDGFRβ (green) co-localization. **C.** Total quantification of PDGFRα volume shows no change in fibroblasts in Control and *Rgs5^ΔPC^* hearts (W4, p=0.4; W8, p=0.39). **D.** Quantification of PDGFRβ total volume increased in *Rgs5^ΔPC^* samples. **E.** Non pericyte-PDGFRβ volume is significantly increased in *Rgs5^ΔPC^* 4 and 8 weeks after tamoxifen injection. **F.** Quantification of PDGFRα^+^ PDGFRβ^+^ volume shows an increase in co-localized cells in *Rgs5^ΔPC^* 4 week after tamoxifen injection. This value is normalized again at 8 weeks (p=0.3). **C-F.** Data are normalized to the vasculature area and expressed as FC from Control. Data are represented as mean ± S.E.M and P values are calculated using two tailed unpaired *t*-test (Control vs *Rgs5^ΔPC^* at W4, n=5; Control and *Rgs5^ΔPC^* at W8, n=8).

